# A single catalytic domain residue regulates the substrate specificity of a mucin-type O-glycosyltransferase in salivary gland function

**DOI:** 10.64898/2025.12.22.695686

**Authors:** Pranav Kumar, Navid Tahan Zadeh, Zulfeqhar A. Syed, Hayley M. Reynolds, Duy T. Tran, Liping Zhang, Kelly G. Ten Hagen, Nadine L. Samara

**Affiliations:** Structural Biochemistry Unit, National Institute of Dental and Craniofacial Research, NIH, Bethesda, MD, 20892, USA; Developmental Glycobiology Section, National Institute of Dental and Craniofacial Research, NIH, Bethesda, MD, 20892, USA; Electron Microscopy Core, National Heart, Lung and Blood Institute, NIH, Bethesda, MD, 20892, USA; University of Utah, School of Medicine, Salt Lake City, UT, 84132, USA; National Eye Institute, NIH, Bethesda, MD, 20892, USA

## Abstract

Human GalNAc-T1 is a ubiquitously expressed mucin-type O-glycosyltransferase that modulates various biological pathways, including immunity, extracellular matrix formation, and salivary gland development and function. However, substrates of GalNAc-T1 are mostly unknown and the mechanistic details of regulation are not well characterized. Here, we investigate the role of the *Drosophila melanogaster* GalNAc-T1 ortholog PGANT5 in salivary glands and provide structural and biochemical details into its mechanism of O-glycosylation. We find that loss of *pgant5* causes irregular secretory granule morphology, altered mucin packaging, and disruption of secretion. Alternative splicing of *pgant5* yields two isoforms, *pgant5A* and *pgant5B*, with higher expression of *pgant5A* in salivary glands. Rescue with *pgant5A*, but not *pgant5B,* partially restores secretory granule morphology and salivary gland function. A single residue difference within the substrate binding pockets of PGANT5A and PGANT5B increases the *in vitro* activity and specificity of PGANT5A towards salivary gland mucins relative to PGANT5B, corroborating the *in vivo* expression and role of *pgant5A* in salivary glands. Our data additionally provide new insights into a catalytic domain motif that regulates GalNAc-T activity. Overall, our studies illustrate how alternative splicing of an O-glycosyltransferase that results in modest amino acid changes can influence substrate specificity to regulate tissue-specific biological functions.

## Introduction

Salivary glands produce and secrete saliva that maintains oral health by preventing infection, lubricating the oral cavity, and assisting with digestion^1^. The protective and lubricating properties of saliva are influenced by proteins that acquire mucin-type O-glycans as they pass through the secretory pathway, a process that is initiated by a family of N-acetylgalactosaminyl (GalNAc) transferases (GalNAc-Ts)^2,3^. Mice lacking *Galnt1,* the gene that encodes GalNAc-T1, are characterized by the abnormal secretion of basement membrane (BM) components during salivary gland organogenesis, resulting in O-glycosylation-dependent endoplasmic reticulum (ER) stress and abnormal gland development^4^. Additionally, GalNAc-T1 is associated with diseases where saliva production is impaired, including Sjögren’s Syndrome, where downregulation of *GALNT1* is observed in patients with severe disease, possibly contributing to the abnormal glycosylation and dysfunction of the salivary gland mucins MUC5B and MUC7^5,6^.

GalNAc-Ts are Golgi-anchored, single pass type II membrane proteins with two luminal domains that are important for activity. The N-terminal catalytic domain adopts a GT-A fold (CAZy family GT-27) with a DHH catalytic motif that coordinates an essential Mn^2+^ and a catalytic flexible gating loop that is involved in substrate binding and alignment (Fig. 1a)^7^. Binding of the donor substrate uridine diphosphate N-acetylgalactosamine (UDP-GalNAc) results in initial loop closure, which is followed by acceptor protein binding and alignment for transfer of GalNAc from UDP-GalNAc to Thr or Ser (Fig. 1a)^8,9^. A short linker joins the catalytic domain to the Ricin B-type lectin domain^10,11^ (carbohydrate-binding module group 13 in the CAZy database) consisting of three repeats termed α, β, and γ, where α and β have the potential to bind substrates by interacting with a prior GalNAc^12^. In humans, 20 isoenzymes are differentially expressed across cell types and regulate diverse biological processes (Fig. 1b)^2,13,14^. Within each cell-type, several GalNAc-Ts are present with distinct and shared acceptor sites on numerous substrates. GalNAc-Ts do not recognize a consensus sequence or motif. Instead, they share ∼35-80% sequence identity within and across species, and each isoenzyme follows its own set of rules for substrate and acceptor site specificity that are encoded within non-conserved residues or motifs^3^.

**Fig 1.**
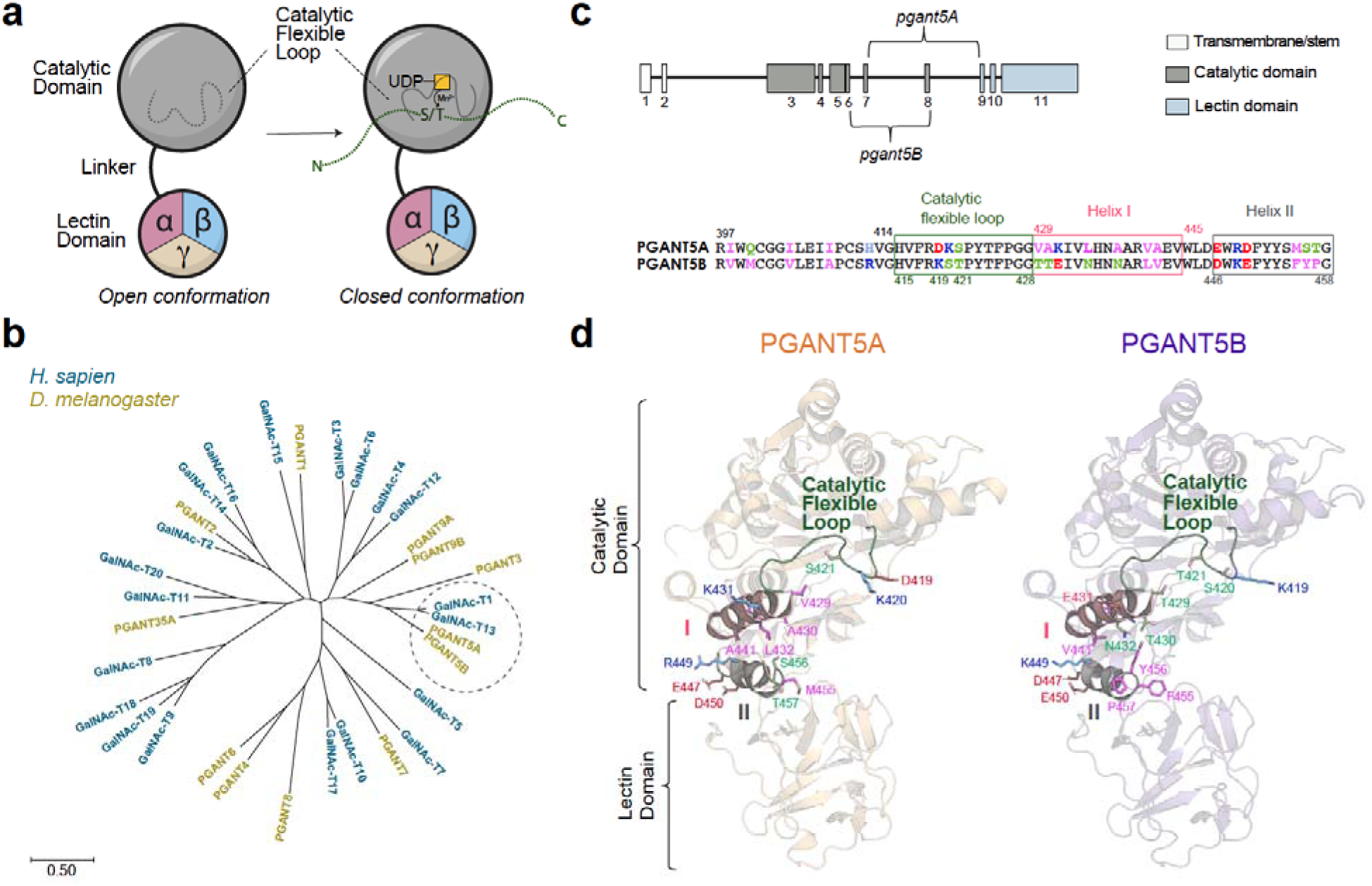
*pgant5* (PGANT5) is the *Drosophila* ortholog of human *GALNT1* (GalNAc-T1) and has 2 splicing isoforms. **a**, GalNAc-Ts contain a catalytic and lectin domain that regulate activity. UDP-GalNAc binding results in the closure of the catalytic flexible loop followed by acceptor peptide substrate binding. The lectin domain contains 3 repeats, α, β, and γ, where α and β, but not γ, can potentially bind GalNAc. **b**, A phylogenetic tree constructed using MEGAX^36^ showing that PGANT5A and 5B are the closest *Drosophila* isoenzymes to human GalNAc-T1. Scale bar represents 0.5 nucleotide substitutions per site. **c**, Alternative splicing of the *pgant5* gene results in *pgant5A* and *pgant5B* that produce distinct isoforms PGANT5A and PGANT5B. A sequence alignment of the spliced region shows that these isoforms differ within the catalytic flexible loop and an adjacent helix (I)-turn-helix (II) motif in the catalytic domain. Hydrophobic residues are magenta, polar residues are light green, basic residues are blue, and acidic residues are red. **d,** Divergent residues are mapped onto AlphaFold2^25^ models of PGANT5A (orange) and PGANT5B (purple) and labeled using the same scheme as **c**.

Despite the central role of GalNAc-T1 mediated mucin-type O-glycosylation in salivary glands, its *in vivo* substrates and regulatory mechanisms have been challenging to characterize due to the complex milieu of GalNAc-Ts and substrates within each cell type. To gain mechanistic insights into mucin-type O-glycosylation and regulated secretion, we use robust genetic tools to investigate these conserved processes in *Drosophila melanogaster (D. melanogaster)*, which has 10 *pgant* genes that encode functional GalNAc-Ts (Fig. 1b, PGANT)^15^. The closest ortholog to GalNAc-T1 is PGANT5 (∼57% identical; Fig. 1b), which is expressed in salivary glands from embryonic to larval stages and has similar substrate preferences to GalNAc-T1^16–18^. Ubiquitous RNAi to *pgant5* results in complete loss of viability^18^, and tissue specific RNAi to *pgant5* in the digestive system results in midguts with a basic pH, revealing a role in regulating gut acidification^18^. However, the functions of *pgant5*/PGANT5 within salivary glands are not known. Interestingly, there are two *pgant5* alternative splicing isoforms termed A and B (UniProt database entries Q6WV17-1 and Q6WV17-2) that differ within residues 398-457 containing the catalytic flexible gating loop and a conserved helix (I)-turn-helix (II) motif of unknown function (Fig. 1c and d).

Here, we investigate the expression and function of *pgant5* in salivary glands and identify how amino acid differences between the tissue-specific splice variants of *pgant5* influence substrate specificity. We show that loss of *pgant5* results in abnormal secretory granule formation and secretion and disrupts the O-glycosylation of salivary gland mucins. Moreover, *pgant5A* shows higher expression in salivary glands compared to *pgant5B,* and rescue with *pgant5A* partially restores the aberrant phenotypes, while rescue with *pgant5B* does not. We find that the enzymes PGANT5A and PGANT5B have distinct substrate preferences, and a single residue difference in their catalytic flexible loops increases the relative enzymatic efficiency of PGANT5A towards positively charged peptide substrates from salivary gland mucins. Overall, our findings show that alternative splicing of the GalNAc-T1 ortholog PGANT5 regulates secretory granule morphology and secretion in salivary glands by producing subtle changes between isoforms to increase the specificity of PGANT5A towards salivary gland substrates. These data provide additional insights into GalNAc-T1 function and illustrate how an individual catalytic domain motif (helix I-turn-helix II) influences GalNAc-T substrate binding and specificity.

## Results

### *Pgant5* regulates secretory granule morphology and secretion

To evaluate the role of *pgant5* in salivary gland function, we performed tissue-specific RNAi-mediated knockdowns of *pgant5* using two RNAi lines (*pgant5^RNAi1^* and *pgant5^RNAi2^*) crossed to the c135-Gal4 driver as described previously^19–21^ (Fig. S1a-b). Using a GFP-tagged mucin Sgs3 (Sgs3-GFP) to track secretory granules^19–23^, we observed a decrease in secretory granule circularity upon loss of *pgant5* relative to wild-type (WT) (Fig. 2a and b). Likewise, we also observed defects in the secretion of the Sgs3-GFP mucin upon RNAi to *pgant5*. As shown in Fig. 2c, secreted Sgs3-GFP normally coats the ventral surface of the prepupa, with none being detected in the salivary glands as development proceeds. However, upon RNAi to *pgant5*, approximately 20% of animals failed to secrete the Sgs3-GFP from the salivary glands, indicating defects in secretion (Fig. 2c). Collectively, these data demonstrate that *pgant5* is required for normal secretory granule maturation and secretion.

**Fig. 2.**
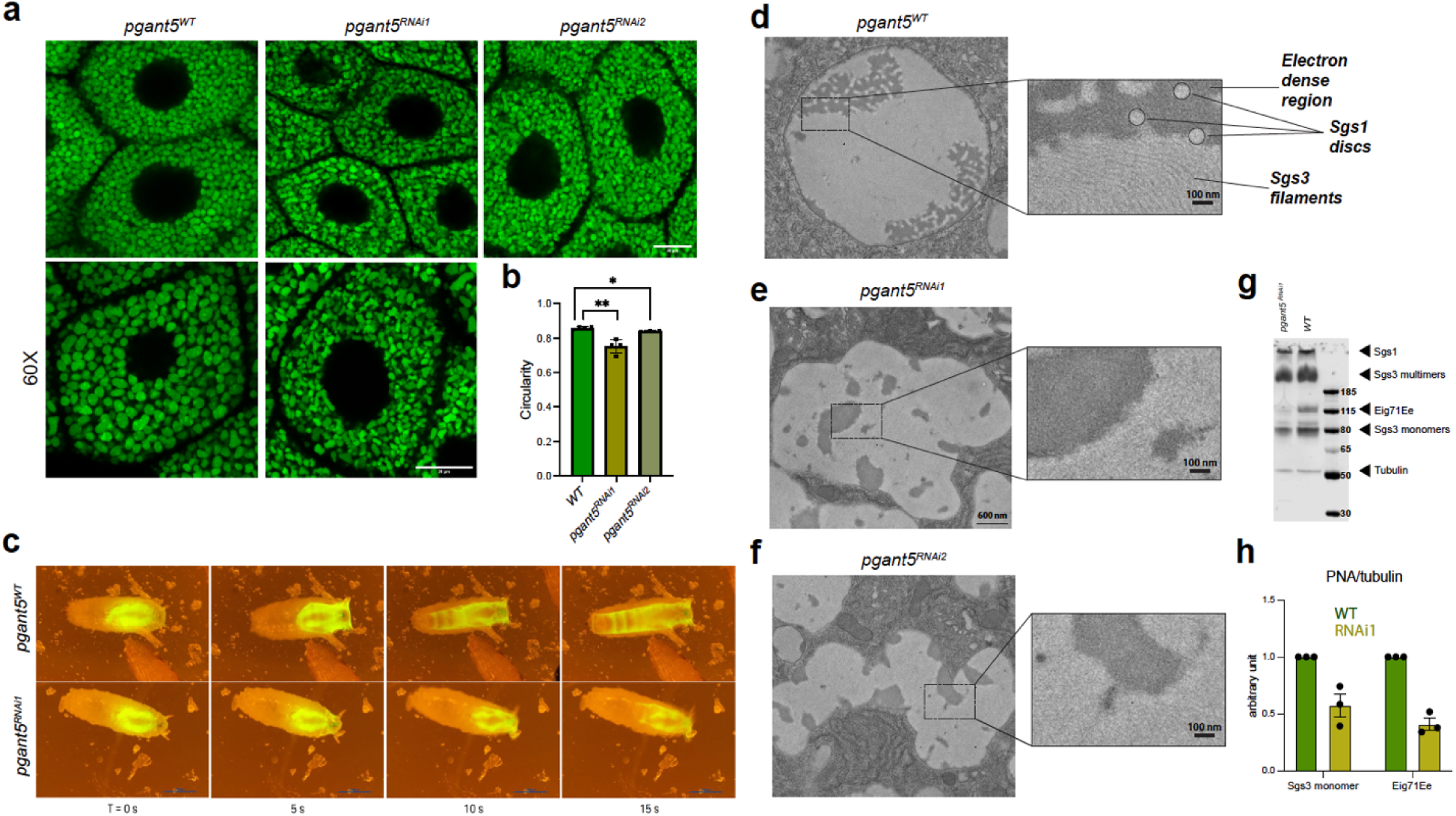
*pgant5* influences salivary gland morphology and secretion. **a,** RNAi knockdown of *pgant5* using 2 RNAi lines as described in the text. Both *pgant5^RNAi1^*and *pgant5^RNAi2^* lines produce secretory granules that have altered morphology relative to *pgant5^WT^*. Scale bars are 20 μM for the top and 20 μM for the bottom 60X magnified images. **b,** circularity measurements showed significant reduction in *pgant5^RNAi1^* and *pgant5^RNAi2^* lines relative to *pgant5^WT^*. **c,** time series images of larval glue expectoration behavior show defective glue secretion in *pgant5^RNAi1^*compared to *pgant5^WT^* over 15 s, * p < 0.05, ** p < 0.01. Transmission electron micrographs (TEM) of a mature secretory granule from a stage 2 *pgant5^WT^* salivary gland, **d**, Magnified views of the boxed regions show that the mature granule has least three distinct structures: Sgs3 bundled filaments, Sgs1 electron-lucent discs, and an electron-dense area. Knockdown of *pgant5 (pgant5^RNAi1^*,**e** or *pgant5^RNAi2^*, **f**) resulted in disorganized Sgs3 bundles and less distinct Sgs1 discs. Boxed areas are shown magnified to the right of each image. Scale bars are 600 nM for images on the left and 100 nm for magnified areas. g, Western blot of *pgant5^WT^* and *pgant5^RNAi1^* salivary glands probed with the lectin PNA and the tubulin antibody. Size markers are shown to the right, as are the positions of the mucins and tubulin control. **h**, quantitation of the ratio of PNA staining/tubulin for Sgs3 monomers and Eig71Ee in *pgant5^WT^* (green bars) and *pgant5^RNAi1^* (yellow bars).

We next used transmission electron microscopy (TEM) to visualize how the absence of *pgant5* may affect normal mucin compaction and restructuring within secretory granules (Fig. 2d-f). As reported previously^21^, mucins within WT secretory granules undergo ordered restructuring to form circular electron lucent discs (Sgs1) or bundles of parallel filaments (Sgs3)^21^ (Fig. 2d). RNAi to *pgant5* results in irregularly shaped granules, with less prominent Sgs1 discs and disordered Sgs3 filaments (Fig. 2e, f). Since both Sgs1 and Sgs3 contain multiple Thr acceptor sites, O-glycosylation by PGANT5 may influence the formation of ultra-structures. Western blotting using a lectin that detects O-linked glycans (Peanut agglutinin; PNA) does indeed show that RNAi to *pgant5* reduces the glycosylation of the mucins Sgs3 and Eig71Ee (Sgs1 is not well resolved on this gel due to its large size and cannot be readily quantified) (Fig. 2g, h), suggesting that at least one role of PGANT5 in salivary gland function is mucin O-glycosylation.

### *pgant5A* overexpression partially rescues knockdown phenotypes

The *pgant5* gene has 2 alternative splicing isoforms that differ within the catalytic domain catalytic flexible loop, and helices I and II (Fig. 1c and d). However, it is not known how these putative isoforms express within salivary glands and if they influence salivary gland function. We used RT-PCR to assess differential expression of *pgant5A* and *pgant5B* and found that *pgant5A* is the predominant salivary glands isoform (Fig. 3a). Thus, we hypothesized that *pgant5A*, but not *pgant5B*, would rescue the RNAi knockdown phenotypes. We constructed rescue lines that overexpressed either *pgant5A* in the RNAi background (*pgant5^RNAi^*, *pgant5A^OE^*, Fig. S1c) or *pgant5B* (*pgant5^RNAi1^, pgant5B^OE^*, Fig. S1d). Granules produced by the *pgant5A* overexpression line appear similar to *pgant5A^WT^*(Fig. 3b-d) while granules produced by the *pgant5B* overexpression line appear similar to *pgant5^RNAi1^*, (Fig. 3b, c, and e) suggesting that *pgant5A* rescues the *pgant5* knockdown phenotypes. Indeed, *pgant5A* overexpression partially restores granule circularity (Fig. 3f) and size (Fig. 3g), and restores secretion (Fig. 3h), while *pgant5B* overexpression does not (Fig. 3f-h). Taken together, the *in vivo* data show that *pgant5A* is the predominant isoform in salivary glands and *pgant5A,* but not *pgant5B,* can partially rescue abnormal secretory granule phenotypes observed upon the loss of both isoforms. To gain insight into why *pgant5A* but not *pgant5B* can rescue the abnormal phenotypes, we next interrogated the glycosylation of salivary gland mucins by the enzymes PGANT5A and PGANT5B *in vitro*.

**Fig. 3.**
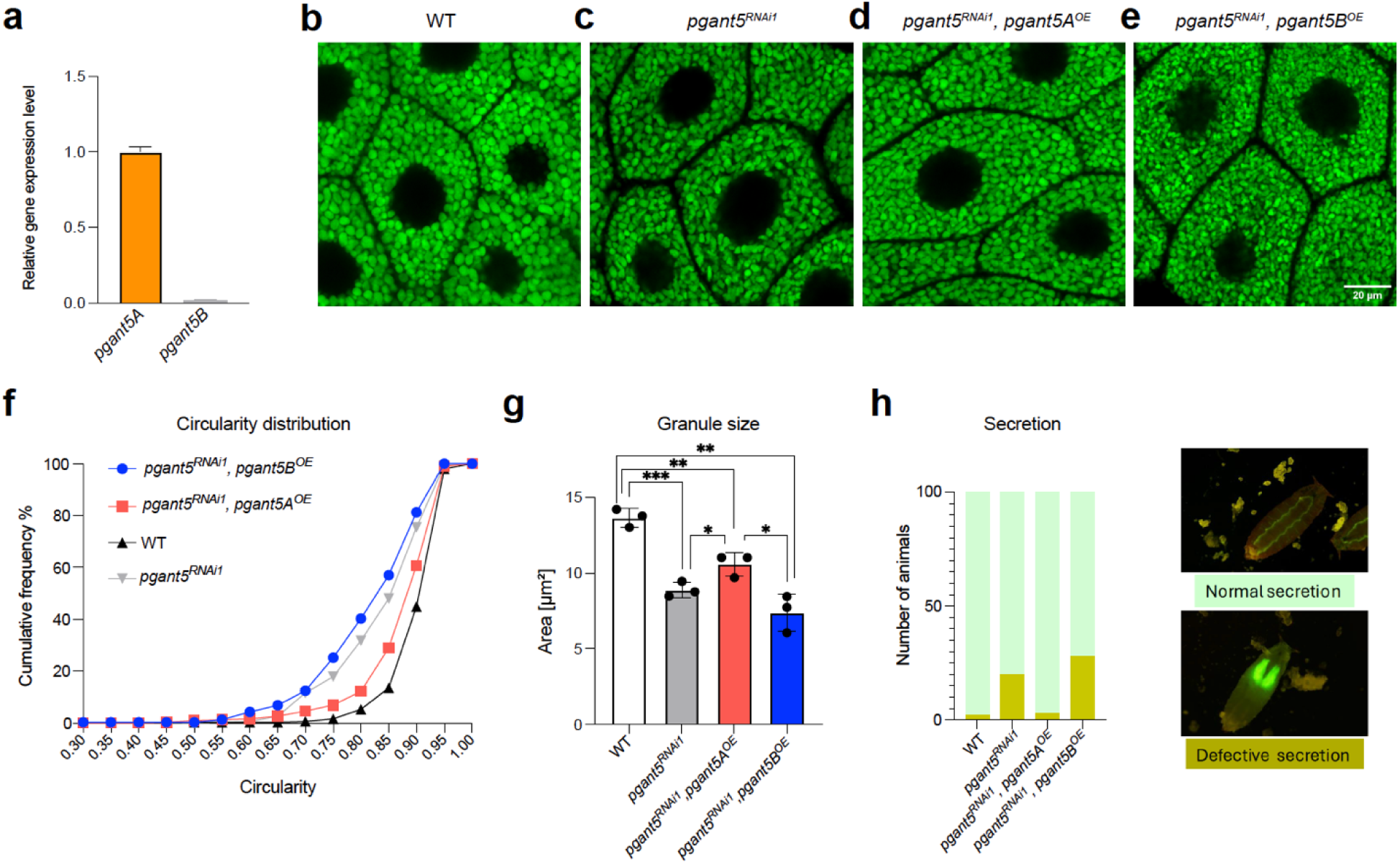
Overexpression of *pgant5A*, but not *pgant5B*, rescues secretion. **a,** qPCR results show that *pgant5A* is the major expressed isoform in the salivary glands. **b-e,** Overexpression of *pgant5A* (*pgant5A*^OE^), but not *pgant5B (pgant5B^OE^*) improves granule circularity and size. **f,** The cumulative circularity distribution graph shows the restoration of granule circularity upon expression of *pgant5A^OE^* in the *pgant5^RNAi1^* background. **g,** granule size comparison among genotypes, and **h,** Quantification of animals with defective glue secretion. Examples of normal and defective secretion are shown to the right. * p < 0.05, ** p < 0.01, *** p < 0.001.

### A catalytic flexible loop residue dictates PGANT5 substrate specificity

To gain mechanistic insight into why *pgant5A*, but not *pgant5B*, partially rescues *pgant5* knock down phenotypes in salivary glands, we first asked if the two isoforms PGANT5A and PGANT5B have distinct O-glycosylation activities against peptide substrates derived from the salivary gland mucins Sgs1 and Sgs3. We show that PGANT5A and PGANT5B have distinct activities towards peptides from the T-rich regions and the mucin domains of Sgs1 and Sgs3 (Fig. S2a, b)^24^. Interestingly, we observe higher initial rates for PGANT5B towards T-rich Sgs1 and Sgs3 substrates (Fig. S2c and d), whereas PGANT5A exhibits higher activity towards the Sgs1 and Sgs3 mucin domain repeat peptides Sgs1-MD and Sgs3-MD (Fig. 4a, S2e). Overall, these initial data show that PGANT5A and PGANT5B have distinct activities *in vitro*.

**Fig. 4.**
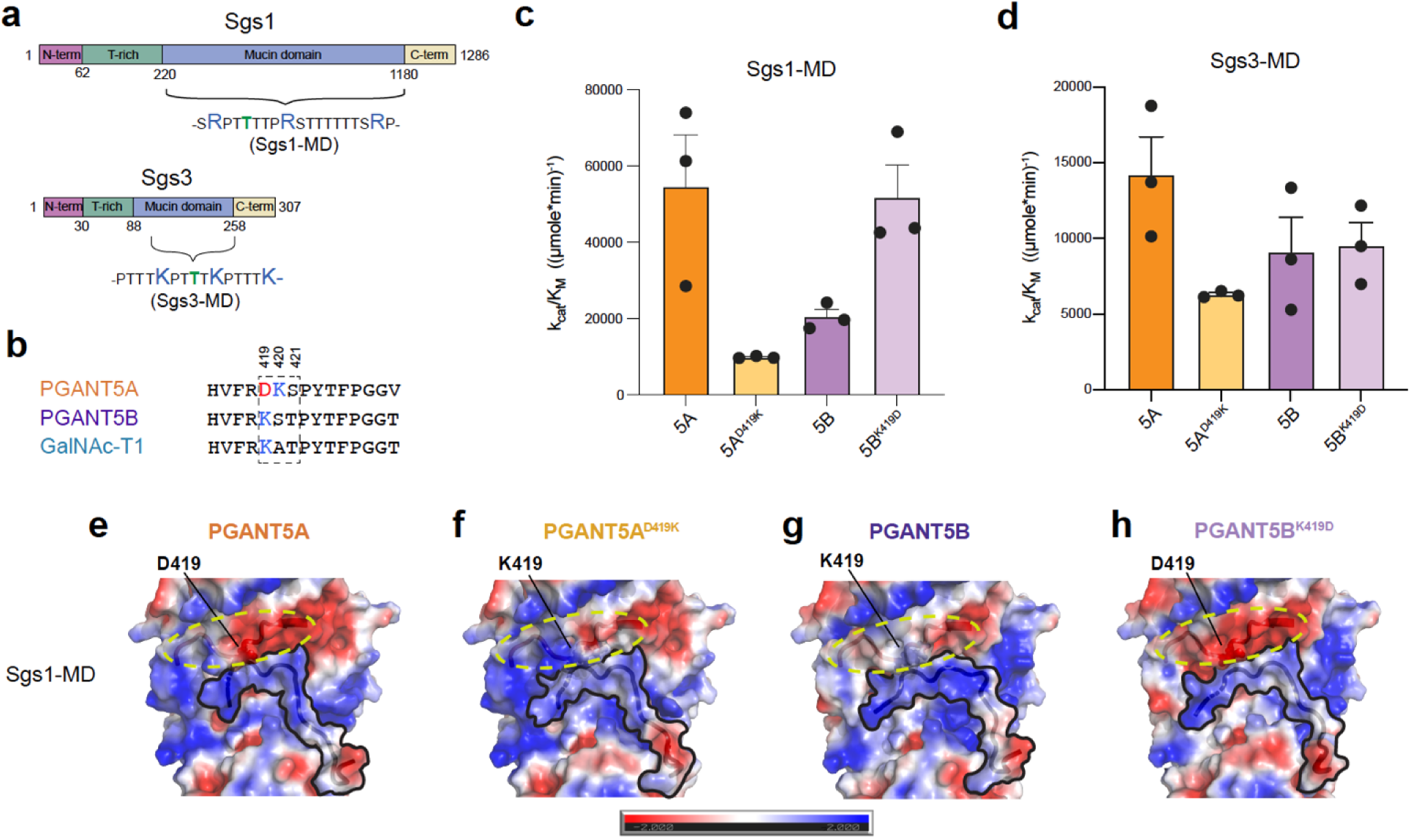
Catalytic loop residue D419 increases PGANT5 activity and specificity towards Sgs1 and Sgs3. **a,** Domain structures of salivary gland mucins Sgs1 and Sgs3. **b,** alignment of the PGANT5A, 5B, and GalNAc-T1 catalytic flexible loops, with residue numbers corresponding to positions in PGANT5. Acidic residues are red, basic residues are blue. **c,** A bar plot of k_cat_/K_M_ showing that PGANT5A glycosylates Sgs1-MD more efficiently than PGANT5B, while activity decreases for PGANT5A^D419K^ relative to PGANT5A^WT^ and increases for PGANT5B^K419D^ relative to PGANT5B^WT^. **d,** PGANT5A is most efficient towards Sgs3-MD, and there is a decrease in activity with PGANT5A^D419K^. Unlike Sgs1-MD, an increase in activity is not observed for PGANT5B^WT^ compared to PGANT5B^K419D^. All data points were derived from experiments done in triplicate 3 different times. Data were analyzed and plotted in GraphPad Prism and the standard errors are shown in the bar graphs. **e-h**, Alphafold2^25^ models of PGANT5A and 5B bound to Sgs1-MD and energy minimized in FlexPepDock^26,27^ showing how the positively charged Sgs1-MD peptides are positioned within the substrate binding groove of PGANT5 variants. The electrostatic surface potentials were calculated with the APBS^28^ electrostatics Pymol plugin using an electrostatic surface potential range from - 2 to +2 for visualization. Red (−2.0) is negative, blue (+2.0) is positive. Figures were generated in Pymol^37^.

To understand the basis of these activity differences, we interrogated divergent residues within the PGANT5A and 5B catalytic flexible loop, which is involved in substrate binding and active site alignment (Fig. 1a, c, and d, Fig. 4b). Residues 419-421 within the catalytic flexible loop are D419, K420, and S421 in PGANT5A, and K419, S420 and T421 in PGANT5B (Fig. 4b). A sequence alignment comparing the spliced region of PGANT5A and 5B to human and *D. melanogaster* GalNAc-Ts (Fig. S3) reveals that only three isoenzymes, including PGANT8, PGANT1, and PGANT5A, have an Asp residue at the position corresponding to PGANT5A aa419, while ∼60% of isoenzymes, including PGANT5B and GalNAc-T1, have a Lys residue at this position. Interestingly, the PGANT5B catalytic flexible loop is identical to the loop in GalNAc-T1, differing by a single aa (S420 in PGANT5B is A349 in GalNAc-T1), whereas PGANT5A and GalNAc-T1 differ at aa419-421 (Fig. 4b). Given that PGANT5A contains a negatively charged Asp residue in the catalytic loop and is highly expressed in salivary glands with proteins containing positively charged mucin domains (pI_Sgs1-MD_=9.95 and pI_Sgs3-MD_=11.61), we hypothesized that D419 enhances the relative specificity and activity of PGANT5A towards these positively charged substrates in comparison to PGANT5B.

To test our hypothesis, we determined the Michaelis-Menten kinetic parameters of the wild-type (WT) enzymes and catalytic loop variants PGANT5A^D419K^ and PGANT5B^K419D^ using the positively charged mucin repeat peptide substrates from Sgs1 and Sgs3 (Fig. 4a). For Sgs1-MD, PGANT5A^WT^ has a ∼2.7-fold higher k_cat_/K_M_ than PGANT5B^WT^ (Fig. 4c), a ∼0.7-fold lower K_M_ and a ∼1.7-fold higher k_cat_ (Fig. 4c, Fig. S4a-c, Table S1). These data suggest that PGANT5A associates with the Sgs1-MD peptide to form a Michaelis (ES) complex and stabilizes the transition state to drive catalysis more efficiently than PGANT5B. A change from D419 to K419 in the catalytic flexible loop of PGANT5A (PGANT5A^D419K^) increases the K_M_ by ∼0.6-fold higher than PGANT5A^WT^ and to a similar value measured for PGANT5B^WT^, indicating that D419 stabilizes substrate binding in comparison to K419. A ∼3-fold decrease in k_cat_ of PGANT5A^D419K^ also points to a catalytic role for D419, resulting in a ∼6-fold decrease in k_cat_/K_M_ compared to PGANT5A^WT^. Interestingly, k_cat_/K_M_ for PGANT5A^D419K^ is 2-fold lower than for PGANT5B^WT^, which could be attributed to the consecutive Lys residues (K419, K420) within the PGANT5A^D419K^ catalytic loop that are more destabilizing for the positively charged Sgs1-MD than the same, less positive region in PGANT5B^WT^ consisting of K419 and S420. In contrast, a change from K419 to D419 in PGANT5B results in a ∼2-fold decrease in K_M_ and ∼1.3-fold increase in k_cat_ resulting a similar k_cat_/K_M_ as PGANT5A^WT^. Overall, the data show how a single acidic amino acid (D419) in PGANT5A increases specificity and activity towards the basic peptide substrate Sgs1-MD.

We observe distinct kinetics for the variants against Sgs3-MD, where the k_cat_ and K_M_ values are ∼5-fold greater and ∼10-fold lower, respectively, than the corresponding Sgs1-MD parameters. This indicates that both the ES and Enzyme-Product (EP) complexes are less stable for the Sgs3-MD peptide than Sgs1-MD, resulting in weaker binding (low on-rate) and higher turnover (high off-rate). Additionally, the differences between PGANT5A and 5B are less pronounced for Sgs3-MD, where PGANT5A has a 1.6-fold greater k_cat_/K_M_ than PGANT5B (Fig. 4d, S4d-f, Table S1). We observed a 2.2-fold decrease in k_cat_/K_M_ for PGANT5A^D419K^ compared to PGANT5A^WT^, and a similar k_cat_/K_M_ for PGANT5B^WT^ and PGANT5B^K419D^. The differences between Sgs1-MD and Sgs3-MD are anticipated given that Arg (Sgs1-MD) is more basic than Lys (Sgs3-MD) and should make stronger interactions with negatively charged residues. Other differences between Sgs1-MD and Sgs3-MD, including length, and position and distribution of charges relative to the acceptor sites could also contribute to variations in activity. How these kinetic parameters translate *in vivo* is further complicated by the extensive mucin domains containing these repeats, with the ∼900 aa Sgs1 mucin domain and the ∼170 aa Sgs3 monomer mucin domain (Fig. 4a). Nevertheless, the data show that PGANT5A has an advantage in glycosylating positively charged mucin repeats, which constitute the largest domains in both Sgs1 and Sgs3. In contrast to the mucin domain peptides, the kinetic parameters for the T-rich substrates Sgs3.3 and Sgs1.4 are similar for both isoforms (Table S2), indicating that the differences between PGANT5A and 5B have no pronounced effects on neutral substrates, and supporting the idea that PGANT5A may have evolved a single negatively charged residue to specifically glycosylate positively charged mucin domains within *Drosophila* salivary glands.

### D419 increases the negative electrostatic surface potential of PGANT5A

To gain insight into the differential glycosylation of peptide substrates by PGANT5A and 5B, we attempted the co-crystallization of PGANT5 in the presence of various peptides but were unsuccessful due to the low yield of enzymes and production of poorly diffracting and irreproducible crystals. We then used AlphaFold2^25^ models of PGANT5A, 5B, and variants, and manually docked peptide substrates into the conserved substrate binding groove, followed by peptide refinement with FlexPepDock^26,27^. Output models were assessed for 2 conserved characteristics: 1) the acceptor Thr is aligned for catalysis in the active site and 2) a Pro residue 3 amino acids C-terminal to the acceptor is properly positioned within a conserved hydrophobic pocket (Fig. 4a, Supplementary dataset 1). APBS^28^ rendered electrostatic surface potential maps show that Sgs1-MD and Sgs3-MD repeat peptides are positively charged within the substrate binding groove, as expected (Fig. 4e-h, Fig. S5a-d). Differences in the electrostatic surface potentials of PGANT5A and 5B occur in multiple regions near the substrate binding groove, including the catalytic flexible loop (yellow dashed circle), which is more electronegative in PGANT5A than PGANT5B due to D419 (Fig. 4e-h, Fig. S5a-d). D419K yields a highly positive version of PGANT5A with two Lys residues in the active site that is not ideal for interacting with positively charged substrates such as Sgs1-MD or Sgs3-MD, explaining why PGANT5A^D419K^ has the lowest k_cat_/K_M_ of the 4 isoenzymes for both peptides (Fig. 4c, d).

Collectively, the data show that a single Asp residue, D419, within the catalytic flexible loop increases the activity of PGANT5 towards positively charged mucin peptides independent of the surrounding charged environment. The increase is due to the formation of a tighter ES complex (lower K_M_) and higher activity (k_cat_). Sgs1-MD is a more optimal substrate than Sgs3-MD for all variants, which could be due to several factors, including amino acid composition, peptide length, and number and position of the acceptor sites, which will dictate how they interact with the various regions of each enzyme to influence binding and activity. The data also show that D419 contributes more to Sgs1-MD than Sgs3-MD specificity based on the larger differences in k_cat_/K_M_ between the D419 variants and K419 variants for Sgs1-MD, indicating that enzymes containing D419 are more sensitive to the charges on Sgs1-MD than Sgs3-MD (Fig. 4c, d).

Indeed, initial rates comparing Sgs1-MD to Sgs1E, where each Arg is substituted for Glu, are higher for Sgs1-MD with enzymes containing D419 (Fig. 5a, b; PGANT5A, PGANT5B^K419D^), and for Sgs1E with enzymes containing K419 (Fig. 5c, d; PGANT5A, PGANT5B^K419D^). We see a distinct trend for Sgs3E, which unexpectedly has a higher rate than Sgs3-MD for PGANT5A, possibly due to higher turnover resulting from a less stable EP complex or because K420 interacts with Glu on this substrate (Fig. S6a). For PGANT5B, both K419 and a nearby R337, which is forming a salt bridge with one of the Glu residues on Sgs3E, are ideally positioned to interact with Sgs3E and increase its activity compared to Sgs3-MD (Fig. S6b, d). It is possible that R337 is also influencing PGANT5A activity towards Sgs3E. Surprisingly, PGANT5A^D419K^ and PGANT5B^K419D^ similarly glycosylate Sgs3-MD and Sgs3E, highlighting the complexity of how these enzymes function (Fig. S6c, d). In all, the data show that PGANT5A has higher specificity than PGANT5B towards positively charged mucin peptides from Sgs1 and Sgs3, but the kinetic basis of the higher specificities is distinct between these two substrates *in vitro*. Nevertheless, the data point to an evolutionary advantage of having higher expression of *pgant5A* relative to *pgant5B* in *Drosophila* salivary glands due to the abundance of positively charged mucins expressed in this organ.

**Fig 5.**
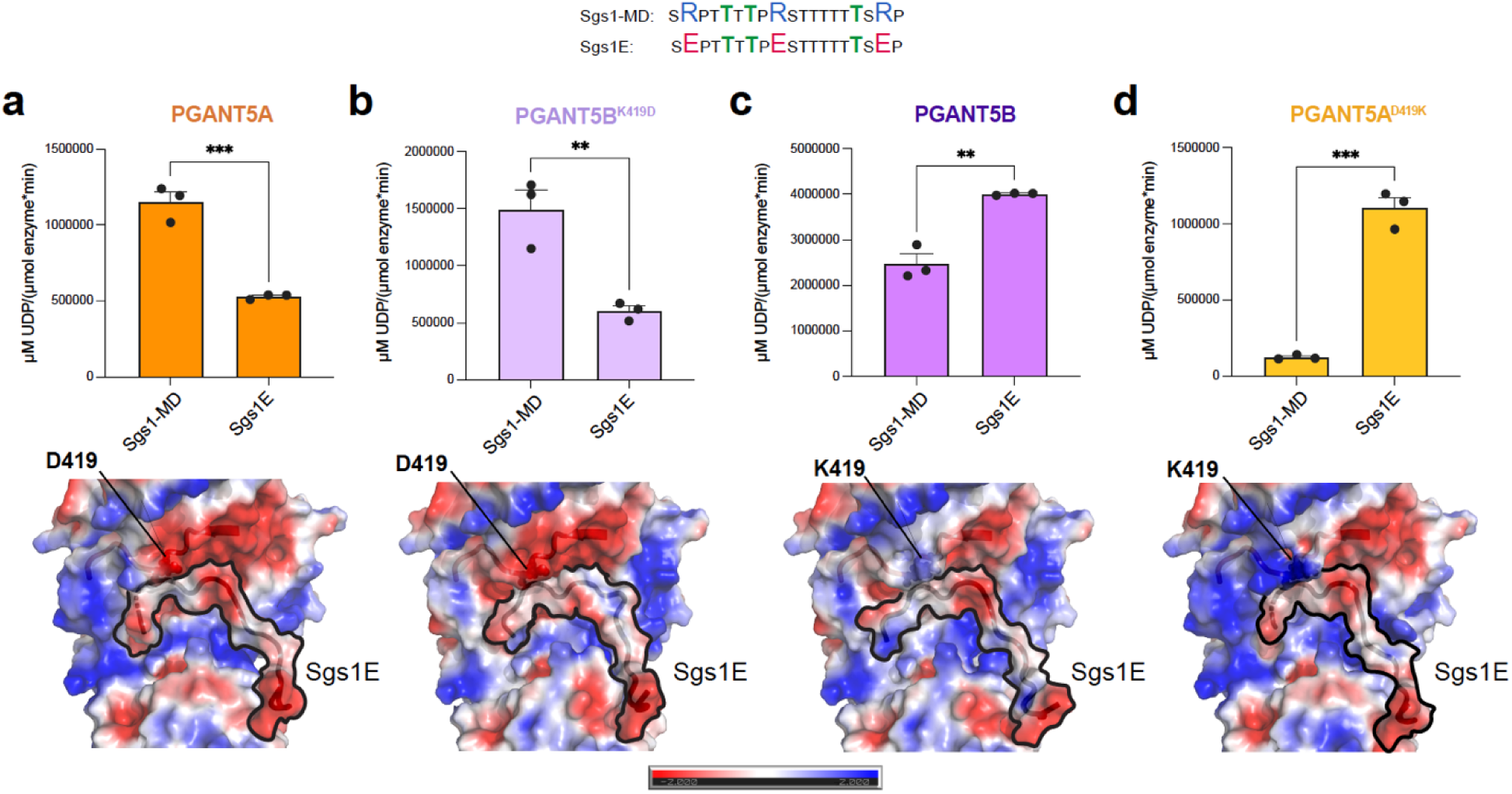
PGANT5A is predominantly responsive to charges on Sgs1-MD. **a-b**, A change from Arg to Glu on Sgs1 peptides (Sgs1E) effects initial rates, where PGANT5A (***p < 0.001) and PGANT5B^K419D^ (**p <0.01) show more activity with Sgs1-MD due to the negatively charged catalytic flexible loop, while **c-d**, PGANT5B (**p < 0.005) and PGANT5A^D419K^ (***p < 0.0005) are more active towards Sgs1-E due to the increased positive charge in the catalytic flexible loop. Corresponding AlphaFold2^25^ models of PGANT5A and 5B bound to Sgs1-MD and energy minimized in FlexPepDock^26,27^ showing how the negatively charged Sgs1E peptides sit within the substrate binding groove of PGANT5 variants. The electrostatic surface potentials were calculated with the APBS^28^ electrostatics Pymol plugin using a electrostatic surface potential range from −2 to +2 for visualization. Red (−2.0) is negative, blue (+2.0) is positive. Figures were generated in Pymol^37^.

### Helix II abuts the lectin domain and influences its function

To assess whether alternatively spliced PGANT5 residues within helices I and II (Fig. 1d) affect PGANT5A and PGANT5B activity, we made single point mutations that changed individual residues from one isoform type to another and purified the enzymes (Fig. S7). The SDS-PAGE (Fig.S7a, b), gel filtration (Fig. S7c, d), and Nano-DSF (Fig. S7e, f) data show that the variants have varied expression levels, but are altogether pure, elute at the same gel-filtration volume, and have comparable T_m_ profiles. Overall, the mutations alter enzymatic activity. However, unlike the catalytic loop residues, these amino acid changes have similar effects on charged mucin repeat substrates and neutral T-rich substrates, suggesting that residues within helices I and II influence enzymatic function independent of substrate amino acid composition (Fig. S8).

Interestingly, single point mutations that change helix-turn-helix PGANT5A residues to PGANT5B residues improve the activity of PGANT5A (Fig. S8). We took advantage of this observation to improve the stability and function of PGANT5A for structural studies, and produced a variant PGANT5A-4M (PGANT5A^A430T, L434N, A437N, S456Y^), which has a higher yield and activity than PGANT5A^WT^, but shows a similar activity trend towards Sgs1 and Sgs3 peptides (Fig. S9a-b). We initially considered using Sgs1-TR as the crystallization substrate because its K_M_ is ∼10 to 20-fold lower than other peptides for both isoforms, and we reasoned that this would promote a tighter enzyme-substrate complex in the crystal (Table S2, Fig. S10a-h). We also designed a substrate based on Sgs1-TR containing GalNAc (Sgs1-TR^GalNAc^) predicting it would interact with the α repeat of the lectin domain and further improve binding. Indeed, a GalNAc on Sgs1-TR^GalNAc^ increases k_cat_/K_M_ by 5-fold for PGANT5A and PGANT5B due to a ∼5-10-fold decrease in K_M_ compared to Sgs1-TR (Table S2, Fig. S10i-p). We disrupted the α repeat GalNAc binding pocket with a D515A mutation that resulted in comparable kinetics to the unglycosylated Sgs1-TR. A change in the β repeat GalNAc binding residue D555A had no effect on K_M_, showing that GalNAc on Sgs1-TR^GalNAc^ likely interacts with the PGANT5 α repeat. Like the WT enzymes, we see activity enhancement for PGANT5A-4M glycosylation of Sgs1-TR^GalNAc^ compared to Sgs1-TR with a 12-fold decrease in K_M_ and 6-fold increase in k_cat_/K_M_ (Table S2, Fig. S10q-u).

We obtained crystals of PGANT5A-4M in the presence of Sgs1-TR^GalNAc^ and solved the structure to around 3.1 Å resolution by fitting and refining four molecules of PGANT5A-4M into the asymmetric unit (Chains A-D, Table S3, Fig. 6a). Although we observed electron density in the substrate binding grooves, it was not sufficient to model the Sgs1-TR^GalNAc^ glycopeptide, because packing of a subset of PGANT5A molecules in the crystal partially occludes substrate binding. Instead, we manually modeled Sgs1-TR^GalNAc^ into PGANT5A-4M by placing an acceptor Thr into the active site and the N-terminal Thr-O-GalNAc into the lectin domain α repeat. Overall, the PGANT5A-4M molecules adopted a similar structure predicted by the AlphaFold2 models, with minor variations. Nevertheless, the crystal structure of PGANT5A-4M provides insight into the dynamics of the alternatively spliced helix-turn-helix region, which has relatively higher B factors than other regions of the catalytic domain in both PGANT5 and GalNAc-T1 (Fig. S11), including the loop that connects the catalytic loop to helix I, and the variant loop region terminating helix II that contains the spliced residues M^455^S^456^T^457^ in PGANT5A and F^455^Y^456^P^457^ in PGANT5B (Fig. 6b).

**Fig 6.**
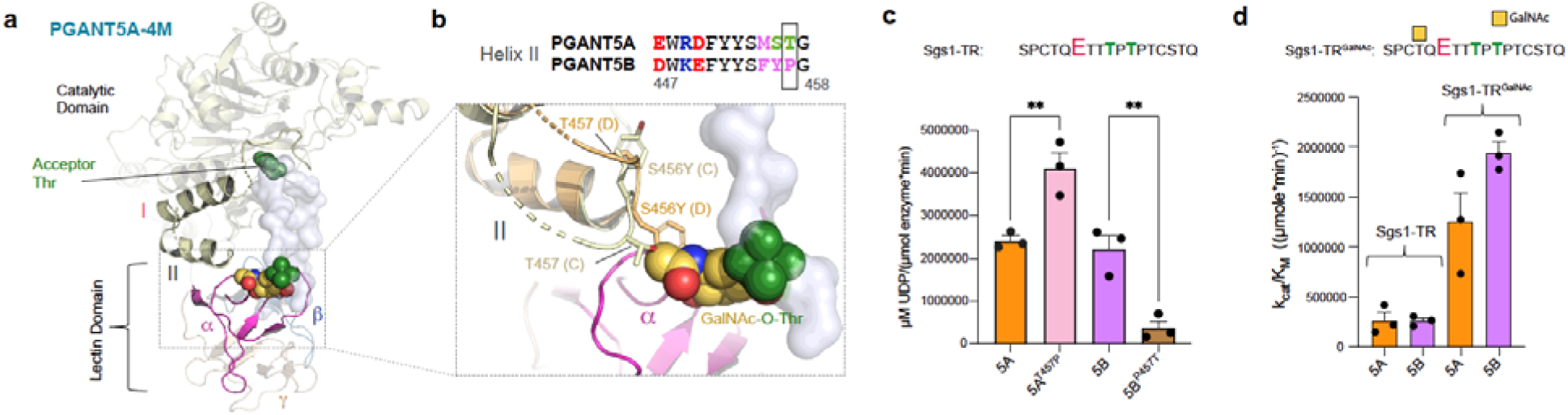
The lectin-catalytic domain interfaces of PGANT5A and PGANT5B are distinct. **a,** Chain C from the X-ray crystal structure of PGANT5A-4M (PGANT5A^A430T, L434N, A437N, S456Y^), solved to 3.1 Å, with Sgs1-TR modeled into the substrate binding groove with the acceptor Thr (green) positioned for catalysis in the active site and Thr-O-GalNAc (green and yellow) positioned in the α repeat, colored magenta. **b,** The loop containing a part of helix II is disordered and residues within that loop overlap with the GalNAc binding pocket (showing the region from chains C in light yellow and chain D in beige), including Y456 (D) and T457 (C). PGANT5A contains a Thr at position 457, while PGANT5B contains Pro, which increases loop rigidity and stability. **c**, PGANT5A^T457P^ has higher activity than PGANT5A^WT^ while PGANT5B^P457T^ is less active than PGANT5B^WT^. (**p <0.05) **d,** While PGANT5A and 5B similarly glycosylate Sgs1-TR peptide, PGANT5B has a higher k_cat/_K_M_ towards a glycosylated version of this peptide Sgs1-TR^GalNAc^, where GalNAc is a yellow square. Activity assays were conducted in triplicate 3 times. Assay data were analyzed and plotted in GraphPad Prism, which was also used to determine the standard error.

The region containing -*MYT-* instead of -*MST-* within each chain of PGANT5A-4M adopts a distinct conformation and is highly disordered and flexible, with weak density for a few residues that were added to the model. For example, in chains D and C, either T457 (chain C) or Y456 (chain D) are positioned near the GalNAc binding pocket within the α repeat of lectin domain, suggesting that these residues could sterically hinder glycopeptide binding (Fig. 6b). The same region in GalNAc-T1 is not disordered in either the apo or substrate bound structure (pdb IDs 1XHB, 8V9Q)^29,30^. Like PGANT5B, GalNAc-T1 contains a Pro residue within this loop (-*IS**P**-*), which may increase structural rigidity and stabilize the region. Indeed, the most pronounced change in helix II occurs in PGANT5B^P457T^, which is considerably less active than PGANT5B^WT^, while PGANT5A^T457P^ is more active than PGANT5A^WT^ (Fig. 6c). The data suggest that a Pro residue at this position could stabilize this region in PGANT5B and GalNAc-T1 to both prevent interference with GalNAc binding and enhance peptide binding, since PGANT5B^P457T^ is also less active than PGANT5B^WT^ against unglycosylated peptides (Fig. S8). In support of more efficient glycopeptide binding by PGANT5B^WT^ relative to PGANT5A^WT^, both PGANT5A and 5B have similar kinetics towards Sgs1-TR, but PGANT5B has higher k_cat_/K_M_ values for Sgs1-TR^GalNAc^ (Fig. 6d).

In all, the data show that the catalytic domain helix-turn-helix motif influences enzymatic activity and modulates lectin domain function through interactions at the catalytic-lectin interface. Whereas PGANT5A has evolved to glycosylate positively charged regions of proteins, PGANT5B and presumably GalNAc-T1 have more efficient lectin domain mediated GalNAc binding resulting in the greater activity enhancement on substrates that contain Thr/Ser-O-GalNAc ∼6-11 amino acids away from the acceptor site. The lectin domain advantage may be more essential to O-glycosylating human salivary gland mucin repeats, which do not contain clusters of positive charges that bind to negatively charged residues or regions of GalNAc-Ts.

### PGANT5 bidirectionality differs from GalNAc-T1

Like GalNAc-T1, both the α and β repeats of PGANT5 contain GalNAc binding pockets. Thus, we asked whether PGANT5 can bind a substrate via α, β, or both repeats simultaneously to position an intermediate acceptor site for catalysis, as was shown for GalNAc-T1^29^. We used a MUC1 (glyco)peptide series to assess how Thr-O-GalNAc at the N-terminus (MUC1-7), C-terminus (MUC1-27) or at both termini (MUC1-7,27) influence activity (Fig. 7a). As shown previously, all glycopeptides enhance GalNAc-T1 activity compared to the unglycosylated peptide (MUC1). Disruption of α repeat GalNAc binding (D444A) primarily decreases MUC1-7 activity, while disruption of β repeat GalNAc binding (D484A) primarily decreases MUC1-27 activity (Fig. 7b)^29^. We observe the same trend for PGANT5A and 5B, suggesting that like GalNAc-T1, PGANT5A and 5B can bind a substrate via α, β, or both repeats simultaneously to enhance activity towards an intermediate site (Fig. 7c-d).

**Fig 7.**
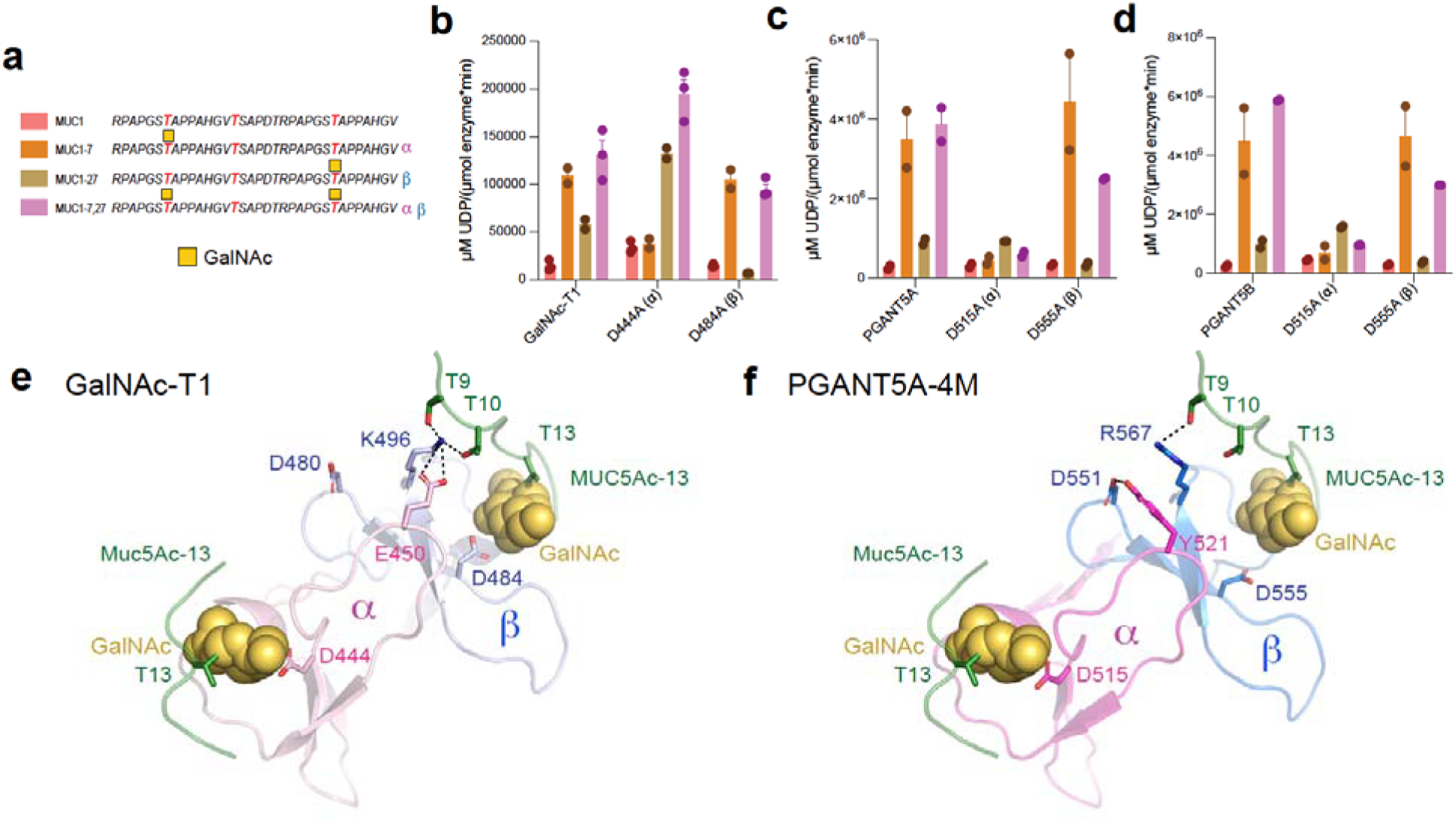
PGANT5A and 5B are bidirectional but α and β function independently. **a,** The MUC1 peptide series used for studying GalNAc-T lectin domain function. Acceptor sites are colored red, and GalNAc is a yellow square. **b,** Initial O-glycosylation rates showing that GalNAc-T1 prefers glycopeptides and uses its α (D444A) repeat to bind GalNAc at the N-terminus and glycosylate in the C-terminal direction (MUC1-7), its β repeat to bind GalNAc at the C-terminus and glycosylate in the N-terminal direction (MUC1-27), and both repeats to glycosylate an intermediate site (MUC1-7,27; **T**SAP). **c and d**, PGANT5A and 5B have similar activity towards MUC1 peptides as GalNAc-T1 with one difference: in GalNAc-T1, disrupting the α repeat (D444A) increases activity towards the β repeat substrate MUC1-27, whereas disrupting the PGANT5 α repeats (D515A) does not affect activity towards MUC1-7, 27. Comparing the lectin domains of **e,** GalNAc-T1 with α in light pink, β in light blue, peptide in green, and GalNAc in yellow to **f,** PGANT5-4M (α in magenta, β in blue) with the MUC5Ac-13 peptide from the GalNAc-T1-MUC5Ac-13 crystal structure superimposed reveals distinct α-β interactions between the two enzymes, where GalNAcT1 E450 on an α loop shared with D444 interacts with K496 on β near the GalNAc binding pocket, an interaction that is not present in PGANT5.

One striking difference between GalNAc-T1 and PGANT5 is that the disruption of α repeat GalNAc binding unexpectedly increases GalNAc-T1 activity towards MUC1-27 and MUC1-7,27, which have a Thr-O-GalNAc at the C-terminus predicted to bind the β repeat. In our previous study, we proposed that the GalNAc-T1 α and β repeats communicate through interface interactions, and disrupting one repeat can change the function of another repeat (Fig. 7b)^29^. However, we do not observe a similar effect for PGANT5, where MUC1-27 is unaffected by an α repeat mutation (D515A), and there is a decrease in activity for MUC1-7,27 due to reduced binding to the N-terminal GalNAc by α (Fig. 7c-d). A comparison of the GalNAc-T1 and PGANT5 lectin domains reveals differences between the interactions at the α-β interface. In GalNAc-T1, the α repeat residue E450 bonds with the β repeat residue K496, which interacts with residues on the peptide substrate and is nearby the β repeat GalNAc binding pocket. A D444A mutation could influence these interactions and influence β repeat function (Fig. 7e). The GalNAc-T1 α repeat residue E450 is Y521 in PGANT5-4M, and it interacts with β repeat residue D551 instead of R567, which is similarly positioned to GalNAc-T1^K496^ (Fig. 7f)^29^. Since D551 is not on the same loop as the β repeat GalNAc binding pocket, interactions between Y251 and D551 may not have the same influence on β repeat function. Overall, these and other possible differences between GalNAc-T1 and PGANT5 within the lectin domain show how even small sequence variations between orthologs can influence function.

## Discussion

This study aimed to understand a branch of GalNAc-Ts that include the ubiquitously expressed, yet still poorly understood isoenzyme GalNAc-T1, which regulates many essential biological pathways. We used *Drosophila*, which contains the GalNAc-T1 ortholog PGANT5, to understand how these family members regulate salivary gland function and secretion. Our studies show that PGANT5 is crucial for normal secretory granule formation and secretion. TEM images of secretory granules indicate that PGANT5 is required for the proper formation of Sgs1 and Sgs3 mucin ultra-structures. Additionally, loss of PGANT5 appears to impact the glycosylation of an additional mucin Eig71Ee. However, these are unlikely the only proteins O-glycosylated by PGANT5, and it is possible that there are other substrates of PGANT5 that influence salivary gland function that have yet to be identified.

Unlike GalNAc-T1, the gene expressing PGANT5, *pgant5*, yields two splicing isoforms, 5A and 5B, that differ within their catalytic flexible loop, a structural feature that is critical for substrate binding and catalysis. *pgant5A*, but not *pgant5B,* is highly expressed in salivary glands and can partially rescue abnormal morphology and secretion phenotypes in *pgant5* knockdowns. Our biochemical studies support a critical *in vivo* role for *pgant5A*, by illustrating that PGANT5A more efficiently O-glycosylates positively charged Sgs1 and Sgs3 mucin repeat peptides than PGANT5B. Intriguingly, the greater specificity of PGANT5A towards these positively charged peptides appears to be dictated by a single residue, D419, within the catalytic flexible loop. Both PGANT5A and 5B can similarly glycosylate neutral regions of the T-rich domains of Sgs1 and Sgs3, suggesting that these regions are negligibly affected by differences in expression or sequence between these two isoenzymes. Collectively, the data provide a model where multiple copies of PGANT5A glycosylate the mucin domains of Sgs1 and Sgs3 like beads on a string, and a few copies of either PGANT5A or 5B can manage the smaller, uncharged T-rich region which contains fewer acceptor sites, (Fig. 8a and b).

**Fig 8.**
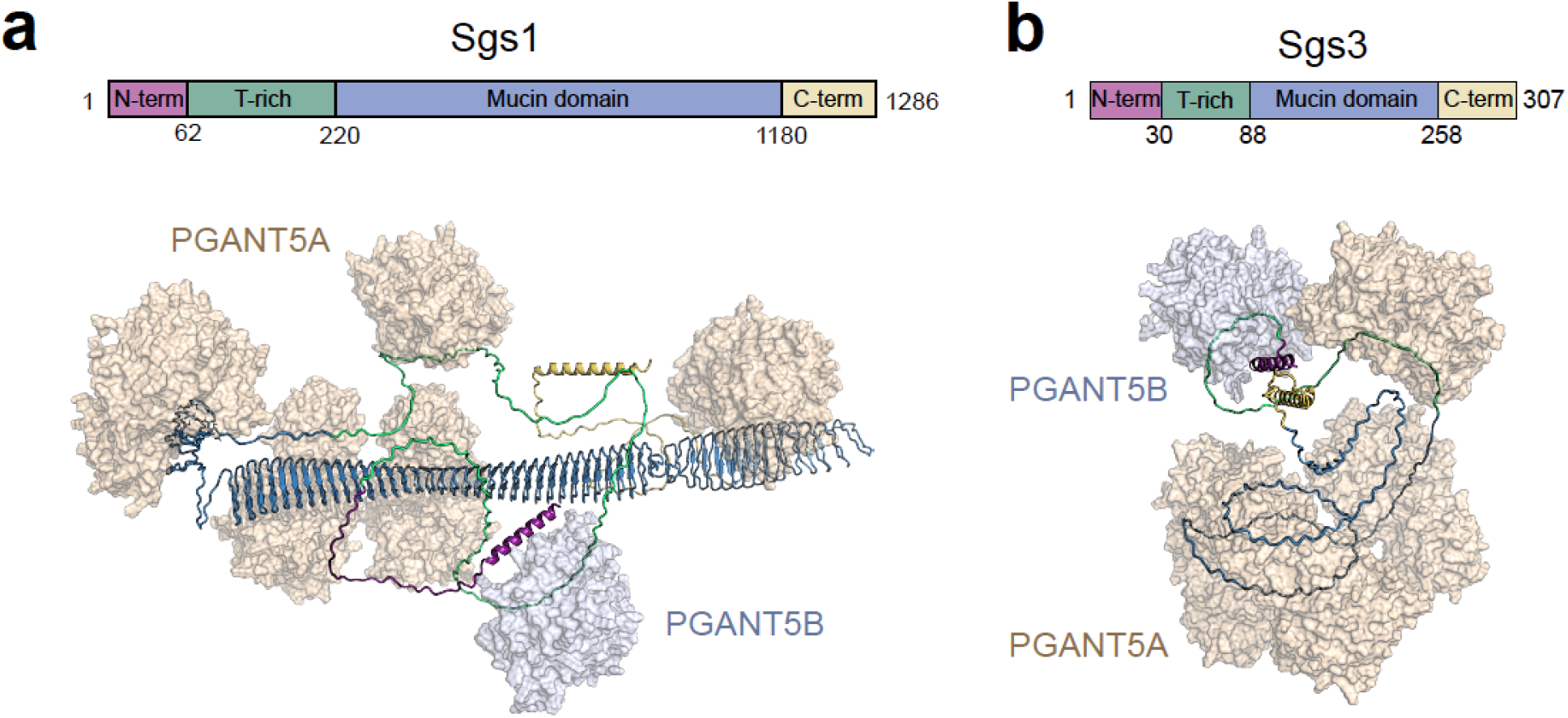
Multiple copies of PGANT5A can glycosylate mucin domains in Sgs1 and Sgs3. **a,** A single Asp residue D419 sufficiently increases the specificity of PGANT5A towards Sgs1, shown here as an AlphaFold2 model with PGANT5A (Beige) molecules bound throughout the mucin domain (blue) like beads on a string. **b,** similar depiction of Sgs3, which contains a smaller mucin domain.

Our data also show that PGANT5A specificity towards Sgs1 is greater than Sgs3, and that D419 has a more important role in binding Sgs1-MD peptides than Sgs3-MD peptides, which turn over more readily as indicated by higher k_cat_ values. This is unsurprising due to the distinct sequences, peptide length, nature of Arg vs Lys, and variations in the positions of charged residues relative to acceptor sites. In the context of a full-length mucin protein, the importance of PGANT5A charge recognition would likely be more crucial due to higher local charges experienced by these enzymes when they dock on the large, repetitive mucin domains *in vivo*. Given that expression of *pgant5A* partially rescues, but does not return everything to WT levels, it is likely that other complexities are at play in the *in vivo* system. There may be as yet other unidentified substrates of PGANT5B or interactions between PGANT5A and 5B that play a role in the overall function of the gland^20^. Nevertheless, the TEM data show that even small changes in O-glycosylation due to the absence of PGANT5 can sufficiently destabilize the formation of Sgs1 and Sgs3 structures. How this is coordinated *in vivo* in the context of other transferases like PGANT9 (which also affects secretory granule morphology)^20^ is unclear, as is the prerequisite for both PGANT5 and PGANT9 for the proper glycosylation of Sgs1 and Sgs3. Given that PGANT5 and PGANT9 share similar *in vitro* activities against these mucin substrates, one would predict redundancy *in vivo*, which is not the case. Clearly, there is much to be learned about the interplay of PGANTS/GALNTs within their native environment and on native mucins.

Interestingly, the GalNAc-T1 catalytic domain is more like PGANT5B than PGANT5A, including within the spliced regions. The catalytic flexible loops of GalNAc-T1 and PGANT5B differ by a single amino acid (Fig.4b), helix I is polar in both GalNAc-T1 and PGANT5B (and hydrophobic in PGANT5A), and the lectin domain adjunct residues within helix II of GalNAc-T1 and PGANT5B contain a Pro residue that is Thr in PGANT5A (Fig. S11a). Since human salivary gland mucins such as MUC1 and MUC7 contain mostly neutral repeats, PGANT5A likely evolved to specifically glycosylate the positively charged salivary gland mucins of *Drosophila*. PGANT5B also has higher specificity towards a glycosylated Sgs1 peptide (Sgs1-TR^GalNAc^) that is predicted to interact with the lectin domain via GalNAc. This may be due to the differences in helix II, where PGANT5B has P457 residue, and PGANT5A has T457 that appears to destabilize this region in the structure of PGANT5A-4M. Although this is not the structure of PGANT5A^WT^, the biochemistry shows that changing T457P increases activity, while P457T decreases activity. Thus, it is likely that PGANT5A evolved to primarily bind substrates in a charge dependent manner, whereas the activities of GalNAc-T1 and possibly PGANT5B are more dependent upon lectin domain GalNAc binding. Additionally, while the GalNAc-T1 α and β repeats communicate to influence each other’s ability to bind substrate, we do not observe a similar result for PGANT5 due to amino acid differences at the PGANT5 α-β repeat interface that result in interactions distinct from those of GalNAc-T1. Essentially, a few amino acid differences between the lectin domains of homologous enzymes influence function, showing how each isoenzyme has evolved to glycosylate mucins within its complex biological milieu.

Indeed, MUC1 or MUC7 mucin repeats do not contain stretches of Thr like Sgs1 and Sgs3 and may require long-range enhancement from lectin domain binding to glycosylate sites that are sparser and more spread throughout the mucin domain compared to Sgs1 and Sgs3. Additionally, these proteins contain a few negatively charged amino acids like Asp, as seen in MUC7 (MUC7: APP**E**TTAAPPT). Another putative GalNAc-T1 substrate is Calnexin which contains Asp near a putative site (SKP**D**TTAPPSS), and MIA3, which contains putative sites surrounded by negative charges that may interact more efficiently with enzymes containing positively charged residues in their catalytic flexible loop, such as GalNAc-T1 and PGANT5B (**DEE**L**D**T**E**YYAV, PLLTFT**D**G**ED**M)^4^. Given that both GalNAc-T1 and PGANT5B have similar catalytic flexible loops, we predict that PGANT5B would more readily glycosylate Tango1, the *Drosophila* homologue of MIA3, than PGANT5A, as well as other substrates with negatively charged residues around the acceptor sites. In conclusion, our study reveals the role of the GalNAc-T1 homologue, PGANT5, in salivary gland function, and our data show that this enzyme is important for mucin structure, secretory granule formation, and secretion, providing insight into how GalNAc-T1 might regulate salivary gland development and function in mammalian systems. Our study also shows how small changes between similar enzymes can have a significant impact on function, whether it is a single Asp residue in the catalytic domain, a Pro at the lectin-catalytic interface, or small variations in the residues at the α-β lectin interface.

## Materials and Methods

### Fly stocks and genetics

All fly stocks and crosses were kept on MM media (KD Medical, Inc.) at 25 °C. The c135-Gal4-driver line (Bloomington #6978) recombined with Sgs3-GFP (Bloomington #5885) was used to induce RNAi in the salivary gland and visualize secretory granules containing Sgs3. For wild type control, w1118 (VDRC#60000) stock was crossed with Sgs3-GFP, c135-Gal4 stock. To induce *pgant5* RNAi, two fly lines from Bloomington Drosophila Stock Center (*pgant5^RNAi1^*: Bloomington #64908, *pgant5^RNAi2^*: #50565) were used. For overexpression of *pgant5A* or *pgant5B* in *Drosophila*, the *pgant5A* and *B* cDNA were cloned into the EcoRI and NotI sites of the pUAST vector to generate *UAS*-*pgant5A*^OE^ and *UAS*-*pgant5B*^OE^. All transgenic lines were generated by BestGene (Chino Hills, CA) using pUAST-*pgant5A*^OE^ and pUAST-*pgant5B*^OE^ vectors. To verify the fly line of *pgant5*^OE^, the forward primer (TCAACTGCAACTACTGAAATCTGCC) and the reverse primer (CTCCACGTCGTATTCCCCG) were used. For rescue experiments, the *pgant5^RNAi1^* line (Bloomington #64908) was recombined with either UAS-*pgant5A*^OE^ or *pgant5B*^OE^, and crossed to the *c135-Gal4, Sgs3-GFP* line.

### Salivary gland imaging and granule circularity measurements

The salivary glands were dissected in Schneider medium and transferred to glass bottom dishes (MatTek) containing 15 μl of medium. A polycarbonate membrane (STERLITECH) was placed on the sample and 15 μl of medium was added on top. A Nikon A1R+ confocal microscope with a Plan Apo IR X 40/1.2 numerical aperture (NA) water immersion (WI) objective was used to image the glands. Circularity measurements were done with Nikon imaging software (NIS-Elements AR version 4.5): 100 individual granules were selected manually, and the circularity was measured using the measurement analysis tool which calculates the circularity based on 4 × π × area/perimeter formula, ascribing 1 to perfect circular granules and ∼0 to elongated polygons. Three biological replicates were used, each with about 100 granules. The average circularity and standard deviation provided by the software were transferred to GraphPad Prism for statistical analysis.

### Salivary gland dissection and Li-COR western blotting

Around 8-10 salivary glands from wandering third instar larvae were dissected and transferred to a 1.5 ml Eppendorf tube containing 50 μl RIPA buffer (Sigma) supplemented with 1 X Halt Protease Inhibitor (Thermo Scientific), followed by homogenization on ice by sonication. Protein samples from about 3 glands were prepared in a reducing loading buffer for SDS-PAGE using a NuPAGE 4–12% Bis-Tris gel (Invitrogen) that was run for about 90 min at 150 V. The proteins were transferred onto 0.45 μm nitrocellulose membranes (Invitrogen). For Li-COR western blotting, the membranes were blocked with Odyssey Blocking Buffer (PBS-based) (Li-COR) for 1 h at room temperature (RT) and then incubated with IRDye 800CW-conjugated PNA (1:3000) and tubulin (1:1000) or Sgs3 antibody (1:2000) overnight at 4 °C. This was followed by 3 × 10 min washing steps with PBS and 0.1 % Tween-20 (PBST), and then incubation with the secondary antibody (anti-rabbit IgG conjugated with IRDye 680 LT diluted in blocking buffer) at RT for 1 hr in the dark. The membrane was washed 3 × 10 min with PBST and then rinsed with PBS for 5 min, followed by imaging using the LI-COR Odyssey CLx and Image Studio software.

### Quantitative RT-PCR

Quantitative RT-PCR (qPCR) was performed using the following PCR primers (*pgant5A* forward: 5′-AATCGCCGTACACCTTCC-3′, *pgant5A* reverse: 5′-ACTCGTCCAGCCAAACTT-3′, *pgant5B* forward: AATCTGGAGATGTCGTTC, *pgant5B* reverse: GGCATTATTGTGATTGACTA). For tissue specific expression and RNAi efficiency of *pgant5a* and *pgant5b,* nine salivary glands from third instar larvae were dissected in PBS and RNA was isolated using RNAqeous Micro Total RNA kit (Invitrogen). cDNA was synthesized using iScript cDNA Synthesis Kit (Bio-Rad). qPCR was performed on a CFX96 real time PCR thermocycler (Bio-Rad) using the SYBR-Green PCR Master Mix (Bio-Rad). RNA levels were normalized to Rp49.

### Expression and purification

PGANT5A and PGANT5B (amino acids 77–626) were cloned from a *Drosophila melanogaster* cDNA template (GenScript, accession numbers NP_608906.2 and NP_001036338.1) into the pPICZα-A expression vector (Invitrogen) for secreted expression of the proteins in *Pichia pastoris (P. pastoris)*. Mutations in PGANT5A and PGANT5B were introduced via site-directed mutagenesis (Q5^®^ High-Fidelity DNA Polymerase, NEB). For *P. pastoris* strain construction, the plasmids were linearized using the PmeI restriction enzyme and transformed into *P. pastoris* SMD1168 cells (Invitrogen) through electroporation. To express PGANT5A, cells were cultured at 30°C in BMGY medium (containing 2% peptone, 1% yeast extract, 1.34% Yeast Nitrogen Base (YNB), 4 x 10-5% biotin, 1% glycerol, 100 mM potassium phosphate at pH 6.0) with 100 μg/ml Zeocin (Invivogen) to an OD600 of ∼20. Expression was induced by centrifuging the cells (1500 x g, 10 min), resuspending in BMMY medium (replacing 1% glycerol with 0.5% methanol), and incubating PGANT5A WT and variants at 16°C and PGANT5B WT and variants at 20°C for 24 h. Following expression, the culture was centrifuged (1500 x g, 10 min), and the supernatant was collected, filtered, and adjusted to pH 7.5 with 50 mM Tris and 10 mM β-mercaptoethanol (βME). Protein purification was performed at 4°C, where the supernatant was loaded onto a 5 ml HisTrap HP column (Cytiva) pre-equilibrated with buffer A (25 mM Tris, 250 mM NaCl, 10 mM βME, and 5% glycerol, pH 7.5). The protein was eluted using a 10 CV gradient of 50-500 mM imidazole. Peak fractions were pooled and digested with TEV protease (1:20 w/w ratio) while dialyzing against 500 ml of buffer A with 25 mM imidazole at 4°C. The His-tagged TEV protease and any uncleaved protein was removed by loading the sample onto a 1 ml His-Trap HP column (Cytiva), followed by washing with 5 ml of dialysis buffer. The glycerol concentration was then increased to 30%, and the protein was aliquoted, flash-frozen in LN2, and stored at −80°C.

### Crystallization, data collection and processing, structure determination and refinement

PGANT5A-4M was thawed and exchanged into crystallization buffer (25 mM HEPES, 100 mM NaCl, 0.5 mM EDTA, 5% glycerol, and 10 mM βME, pH 7.3) using a Superdex 200 Increase 10/300 GL (Cytiva). The peak fractions were concentrated to ∼10-12 mg/ml with a 10 kDa cut-off Amicon Ultra-4 concentrator (Millipore Sigma). The enzyme-peptide-UDP-Mn²⁺ complex was prepared by combining the protein with 5 mM of the glycopeptide Sgs1-TR^GalNAc^, 5 mM UDP-2-(acetylamino)-4-fluoro-D-galactosamine disodium salt (UDP-GalNAc-F, ChemBind), and 5 mM MnCl₂, resulting in a final protein concentration of ∼6.0 mg/ml. Hanging drop vapor diffusion crystallization was set up by mixing equal volumes (1 μl) of the protein complex and reservoir solution (18% PEG 3350 w/v, 0.1 M HEPES pH 7.2, 20 mM MgCl₂), and equilibrated against a 500 μl reservoir solution. Crystals appeared after four days at 20°C in 24-well plates and were cryoprotected with a solution containing 20% glycerol before being flash-frozen in LN2 for X-ray diffraction analysis. X-ray data collection took place at the Brookhaven National Laboratory (BNL), NSLS-II beamlines (New York). Diffraction data were processed and scaled using HKL2000^31^. Molecular replacement was performed with MolRep (CCP4^32^) using an Alphafold2^25^ generated model of PGANT5A as the search model. Models were manually built in Coot^33^ and further refined in PHENIX^34^. Final models were validated using PROCHECK and MOLPROBITY, and structure figures were generated using PyMOL^35^.

### UDP-Glo assays

Glycosyltransferase activity was measured using the UDP-Glo™ Glycosyltransferase Assay kit (Promega, V6961). A 25 μL reaction was initiated by adding either 10 or 50 nM of purified wild-type and mutant PGANT5A, and PGANT5B to 5 mM ultrapure uridine diphosphate (UDP)-N-acetylgalactosamine (GalNAc) (provided in the kit) and an acceptor peptide (Anaspec, Peptide 2.0) in 5 mM MnCl₂ (Sigma Aldrich) and 1X HEPES buffer (100 mM NaCl, 5 mM β-mercaptoethanol, 25 mM HEPES, pH 7.3). Reactions were assembled in a 96-well plate (Greiner Bio-One, 655073) and incubated at 37°C for either 15 or 30 minutes. The reactions were stopped by adding 25 μL of the UDP detection reagent to each well and incubating at 27°C for 60 minutes using a Synergy Neo2 Hybrid plate reader (BioTek) to measure luminescence. A standard curve of UDP in 5 mM MnCl₂ and 1X HEPES buffer was generated for each measurement to relate luminescence to UDP concentration, providing a direct correlation with glycosyltransferase activity within the 60-minute incubation period. Enzyme kinetics were performed in triplicate at different substrate concentrations (0 μM – 2000 μM) and repeated 2 or 3 times. Kinetic parameters, the value of maximum velocity (V_max_) and Michaelis Constant (K_M_) were obtained from each biological replicate and used to calculate k_cat_ and k_cat_/K_M_. Data were analyzed using Microsoft Excel and GraphPad Prism 10. Standard errors were calculated where appropriate and outliers were identified and eliminated using the ROUT method in GraphPad Prism 10.

### Nano-DSF

PGANT5A and PGANT5B variants were thawed and buffer-exchanged into an assay buffer (25 mM HEPES, 100 mM NaCl, 0.5 mM EDTA, 5% glycerol, 10 mM βME, pH 7.3) using a Superdex 200 Increase 10/300 GL column (Cytiva). Protein concentration was achieved using an Amicon ultra-centrifugal concentrator with a 10 kDa cut-off at 4000 x g (Millipore Sigma). Protein concentrations were then measured using the Pierce™ BCA Protein Assay Kit (ThermoFisher Scientific). Denaturation profiles for both isoforms (PGANT5A and PGANT5B) and their variant proteins were analyzed with the Tycho NT.6 (NanoTemper GmbH, Germany). Approximately 10 μL of protein at a concentration of ∼0.2 mg/mL was loaded into NT.6 glass capillaries, followed by heating from 35°C to 95°C at a rate of 30°C/min. Fluorescence intensities were recorded at 330 nm (F330) and 350 nm (F350), and the ratio (F350/F330) along with their first derivatives (∂F330/∂T, ∂F350/∂T, ∂(F350/F330)/∂T) were obtained. The first derivative (F350/F330) versus temperature (°C) plots were generated using GraphPad Prism.

### Transmission electron microscopy

Salivary gland (SGs) for TEM was prepared as described previously^21^. Briefly, SGs were fixed in a mixture of 2% glutaraldehyde and 4% formaldehyde in 0.1M sodium cacodylate buffer, pH 7.4, for 60 min at RT and overnight at 4°C. After overnight fixation, SGs were washed in 0.1 M Sodium cacodylate buffer, pH 7.4, and post-fixed in 1% OsO_4_ in 0.1 M sodium cacodylate buffer for 60 min on ice. Next, SGs were rinsed and washed in water and en-bloc stained with 1% uranyl acetate overnight at 4°C. The following day SGs were rinsed and washed in water and dehydrated through a graded ethanol series (30%, 50%, 70%, 85%, 90% and 2×100%) followed by propylene oxide. SGs were then infiltrated in a gradient mix of propylene oxide and resin (Embed 812 resin) before being infiltrated with 3 changes of pure resin and embedded in 100% resin and baked at 60°C for 48 hours. Ultrathin sections (70 nm) were cut on an ultramicrotome (Leica EM UT7), and digital micrographs were acquired on JEOL JEM 1200 EXII operating at 80Kv and equipped with AMT XR-60 digital camera.

## Data Availability

Structure coordinates and X-ray diffraction data have been deposited in the Protein Data Bank, www.wwpdb.org (PDB ID code: XXXX). Source data are provided with this paper as a Source Data File.

## Acknowledgements

This research used the NYX beam line of the National Synchrotron Light Source II, a U.S. Department of Energy (DOE) Office of Science User Facility operated for the DOE Office of Science by Brookhaven National Laboratory under Contract No. DE-SC0012704 with the assistance of the staff at SER-CAT consortium, which is supported by its <u>member institutions</u>, equipment grants (S10_RR25528, S10_RR028976 and S10_OD027000) from the National Institutes of Health, and funding from the Georgia Research Alliance. Molecular graphics and analyses performed with UCSF ChimeraX, developed by the Resource for Biocomputing, Visualization, and Informatics at the University of California, San Francisco, with support from NIH P41-GM103311. We thank the Bloomington Stock Center and the Developmental Studies Hybridoma Bank for providing fly stocks, antibodies and other reagents. This work was supported by the Intramural Research Program of the NIDCR, National Institutes of Health Grant 1-ZIA-DE000754-03 (to N.L.S.) and 1-ZIA-DE000713 (to K.G.T.H.). This research was also supported in part by the NIDCR Imaging Core (DE000750-01). The contributions of the NIH authors were made as part of their official duties as NIH federal employees, are in compliance with agency policy requirements, and are considered Works of the United States Government. However, the findings and conclusions presented in this paper are those of the authors and do not necessarily reflect the views of the NIH or the U.S. Department of Health and Human Services.

## Author Contributions

NLS, PK, DTT, HR, LZ, and KGTH conceptualized the project. PK cloned, expressed, and purified PGANT5 variants, and conducted structural and biochemical experiments. DTT made initial *in vivo* observations. NTZ made fly lines and performed *in vivo* experiments. ZS conducted TEM experiments. NLS wrote the paper. PK, NTZ, and ZS added Materials and Methods, and PK, NTZ, KGTH, LZ contributed to figure and paper editing.

