## Supplemental Information for "A single catalytic domain residue regulates the substrate specificity of a mucin-type O-glycosyltransferase in salivary gland function"

**Supplemental Figures**

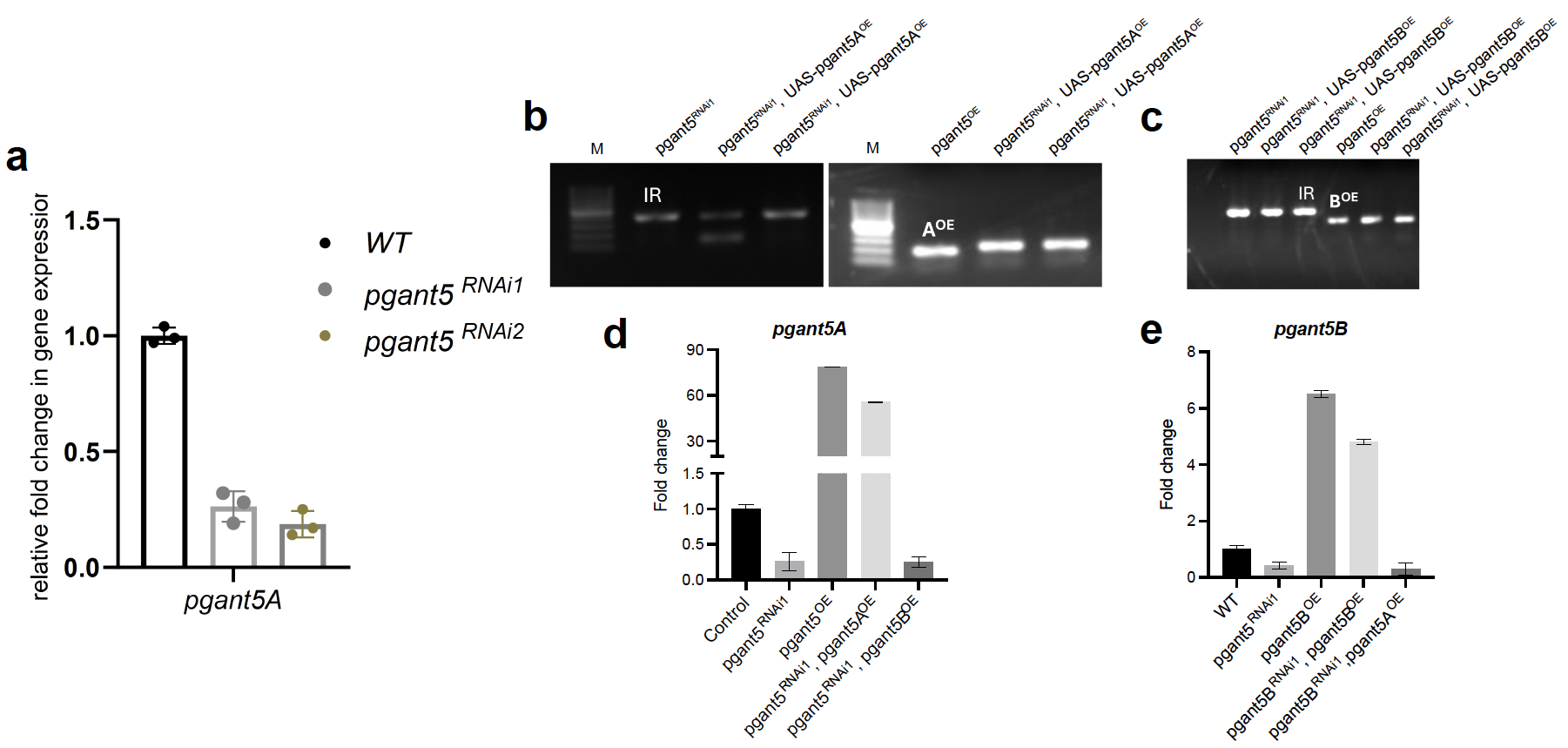

**Supplemental Figure 1. a,** RNAi knockdown lines RNAi1 and RNAi2 resulting in reduced expression of *pgant5A* in the salivary glands as determined by qPCR. **b,** Agarose gel image confirming the rescue construct *UAS-pgant5A^OE^* in the RNAi1 knockdown background and **c,** Agarose gel image confirming the rescue construct *UAS-pgant5B^OE^* in the RNAi1 knockdown background*.* **d,** QPCR of *pgant5A* expression in all the genotypes. **e,** QPCR of *pgant5B* expression in all the genotypes. *M: marker, IR: Inverted repeat, OE: Overexpression*

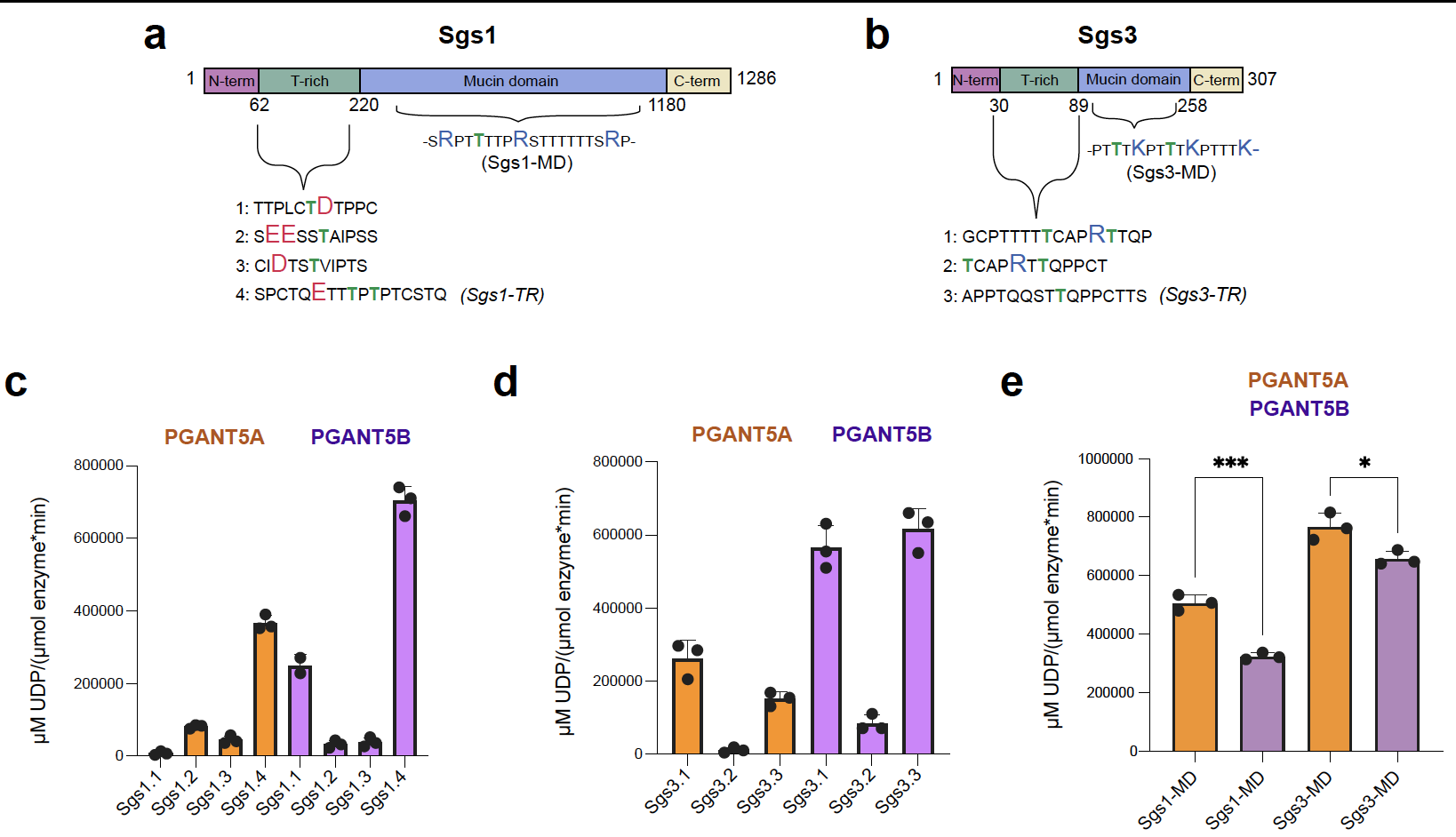

**Supplemental Figure 2. a,** Domain structure of the *Drosophila* mucin Sgs1 and peptides used in this study **b,** Domain structure of the *Drosophila* mucin Sgs3, and peptides used in this study. **c,** Activity of PGANT5A and 5B tested against a panel of Sgs1 T-rich domain peptides. **d,** Activity of PGANT5A and 5B tested against a panel of Sgs3 T-rich domain peptides. **e,** Activity of PGANT5A and 5B tested against mucin domain peptides Sgs1-MD and Sgs3-MD (***p < 0.0005, *p < 0.05).

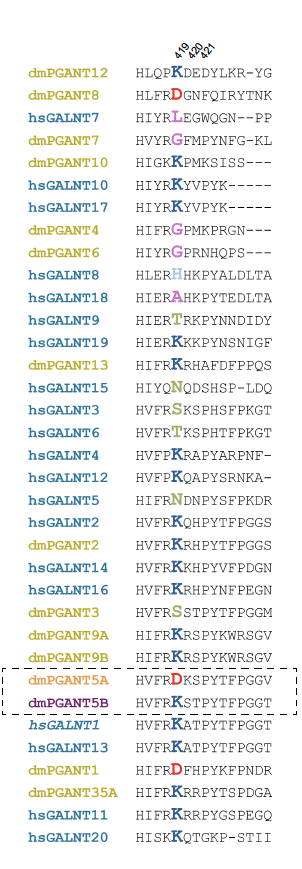

**Supplemental Figure 3.** Catalytic flexible loop sequence alignment of human GalNAc-Ts and *Drosophila* PGANTs.

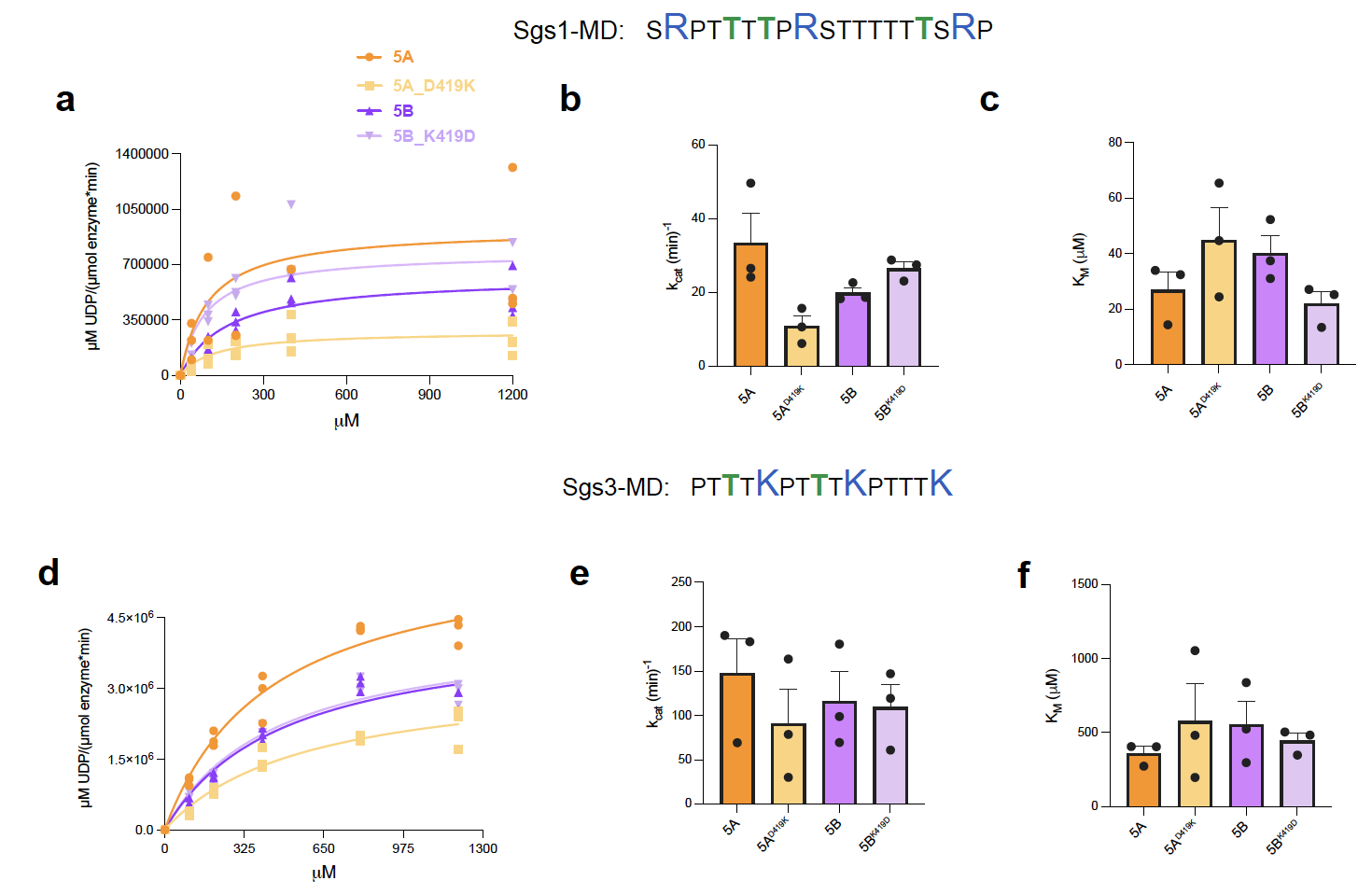

**Supplemental Figure 4. a,** PGANT5 variants kinetics assaying activity towards Sgs1-MD. **b,** A bar plot showing k_cat,_ and **c,** K_M_. **d,** PGANT5 variants kinetics assaying activity towards Sgs3-MD. **e,** A bar plot showing k_cat,_ and **f,** K_M_. All assays were conducted in triplicate 2-3 different times, and data were analyzed in GraphPad Prism as described in Materials and Methods. Data are tabulated in Table S1.

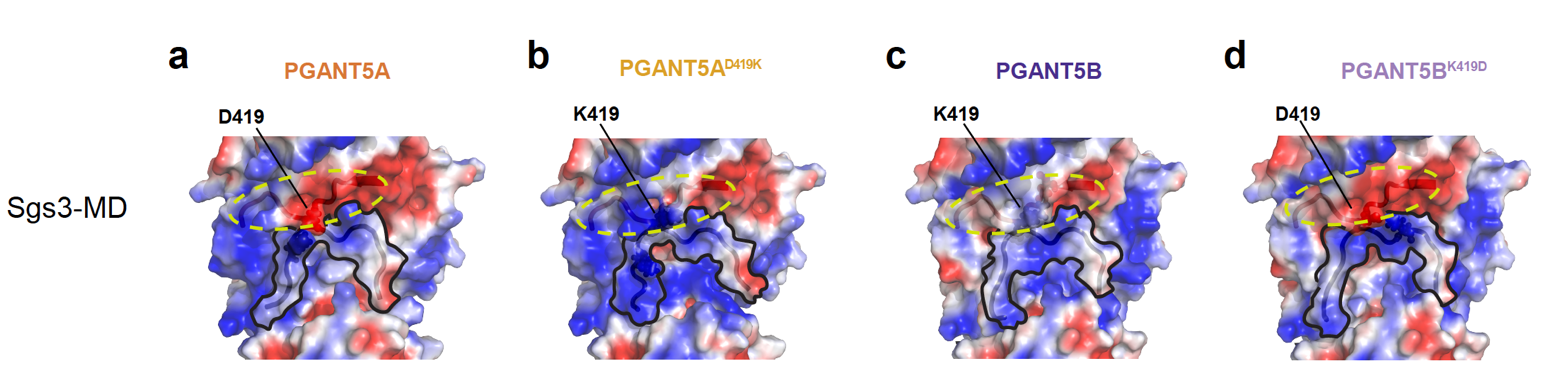

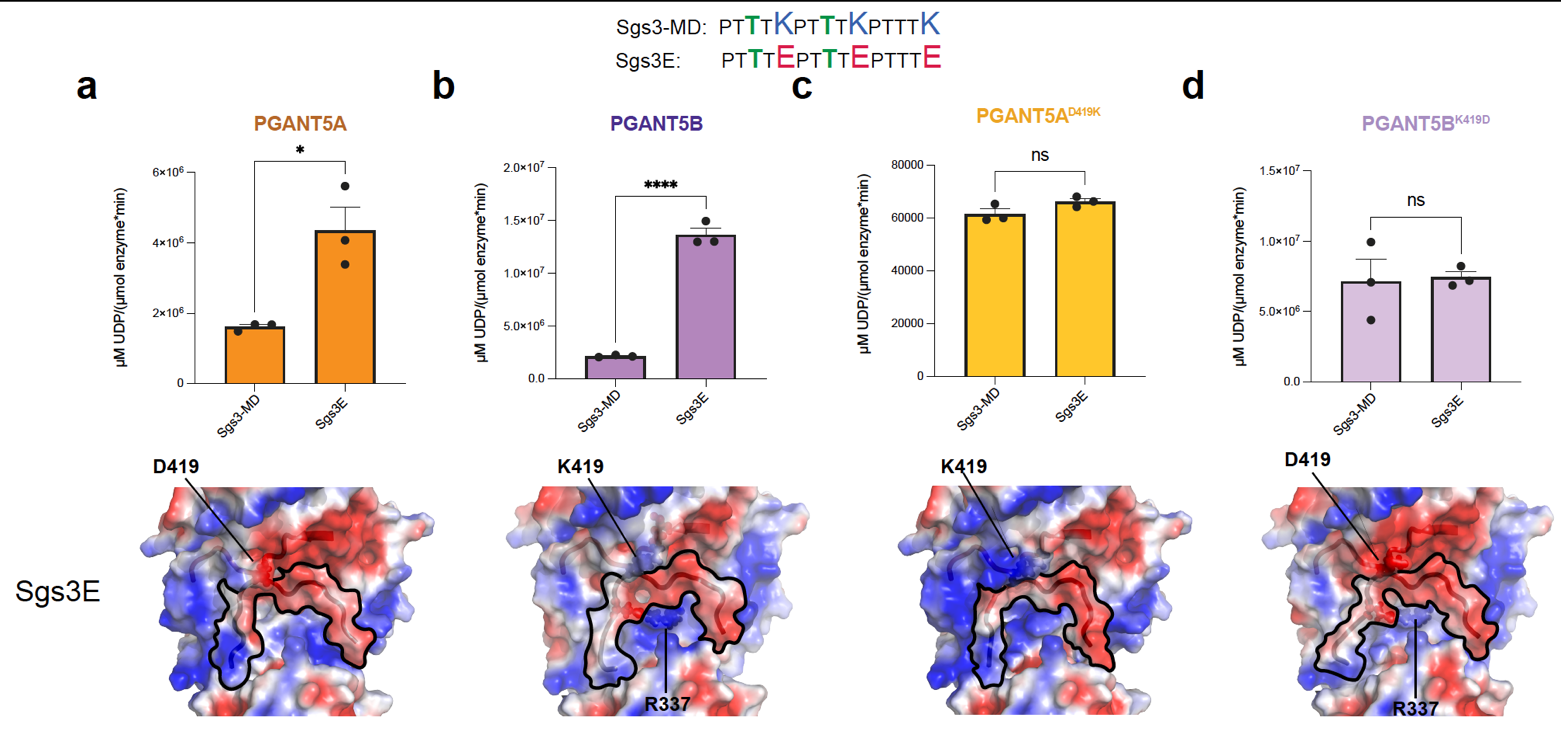
**Supplemental Figure 5. a-d,** AlphaFold2 structures of each variant with Sgs3-MD docked using FlexPepDock. The catalytic flexible loops are encircled with yellow dashes.

**Supplemental Figure 6. a-d,** Assays comparing the activity of PGANT5 variants using Sgs3-MD and its negatively charged counterpart Sgs3E. AlphaFold2 structures of each variant with Sgs3E docked using FlexPepDock (*p < 0.05, ****p < 0.0001).

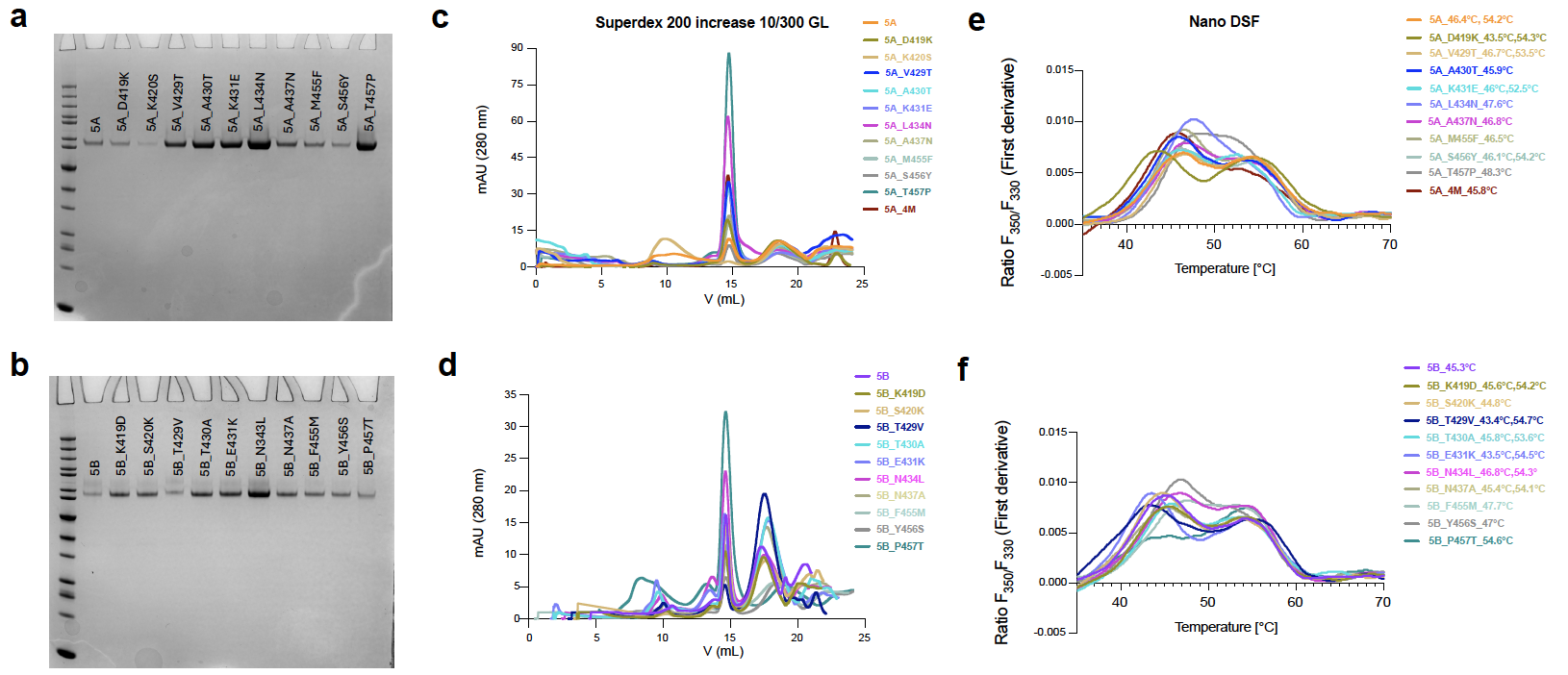

**Supplemental Figure 7. a, b,** SDS-PAGE gels showing purified PGANT5A and 5B variants. **c, d,** Gel filtration analyses of variants using the SD200 Increase 10/300 GL column. **e, f,** Nano-DSF data for PGANT5 variants.

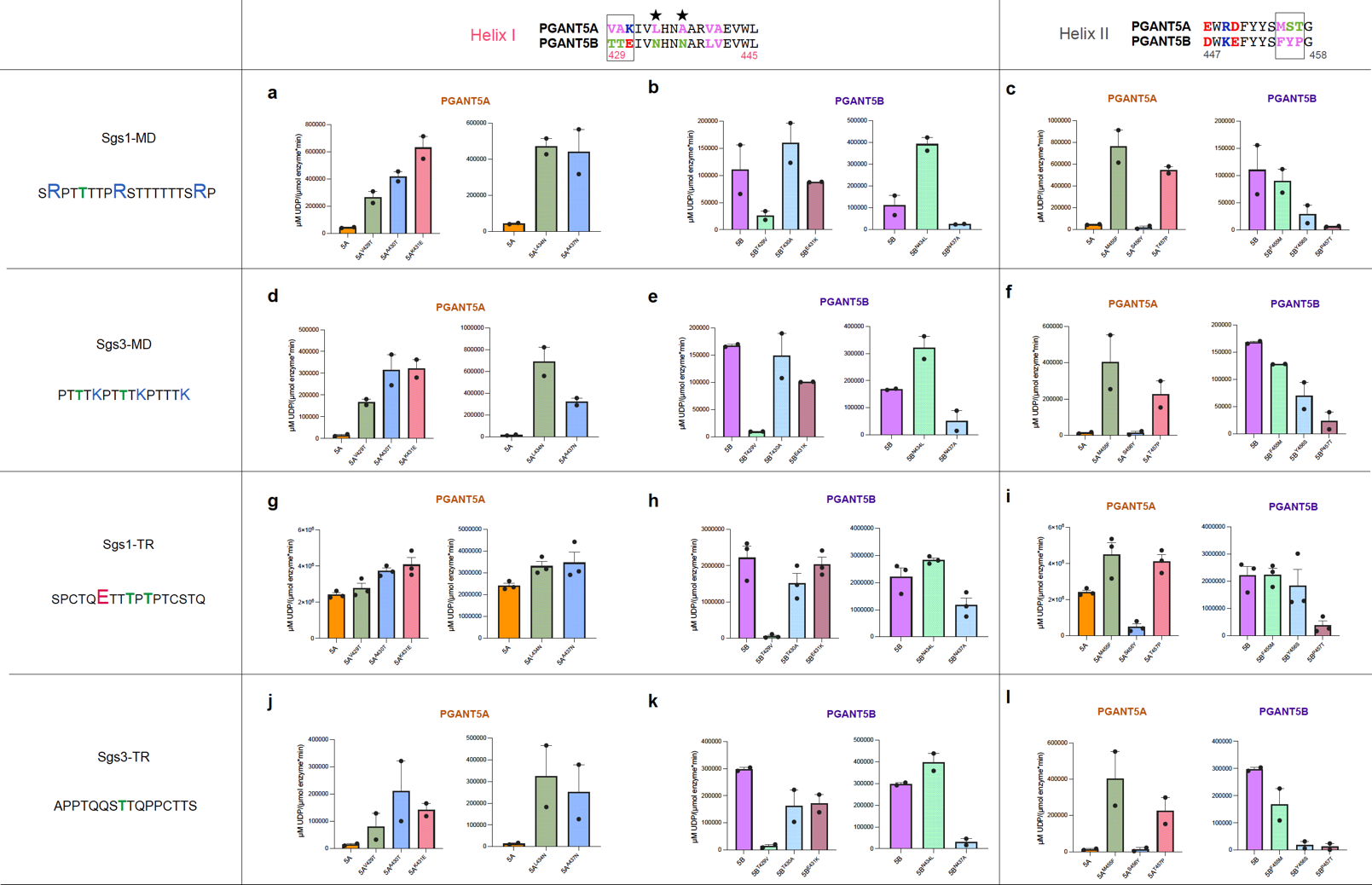

**Supplemental Figure 8.** Activity assays for helix I and II PGANT5 variants using mucin domain (MD) and T-rich (TR) domain substrates.

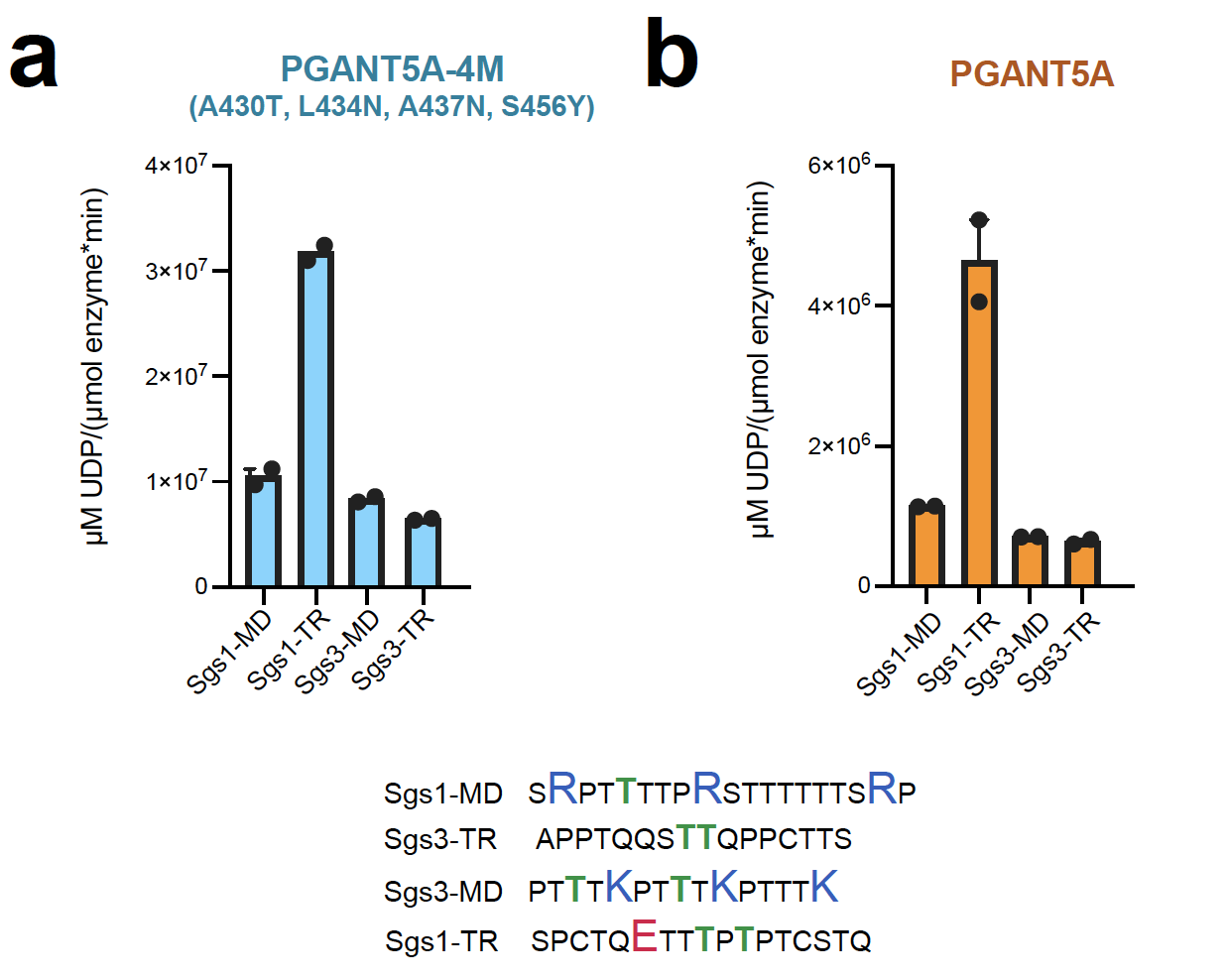

**Supplemental Figure 9.** Activity comparison of PGANT5A-4M and PGANT5-WT using mucin domain (MD) and T-rich (TR) substrates.

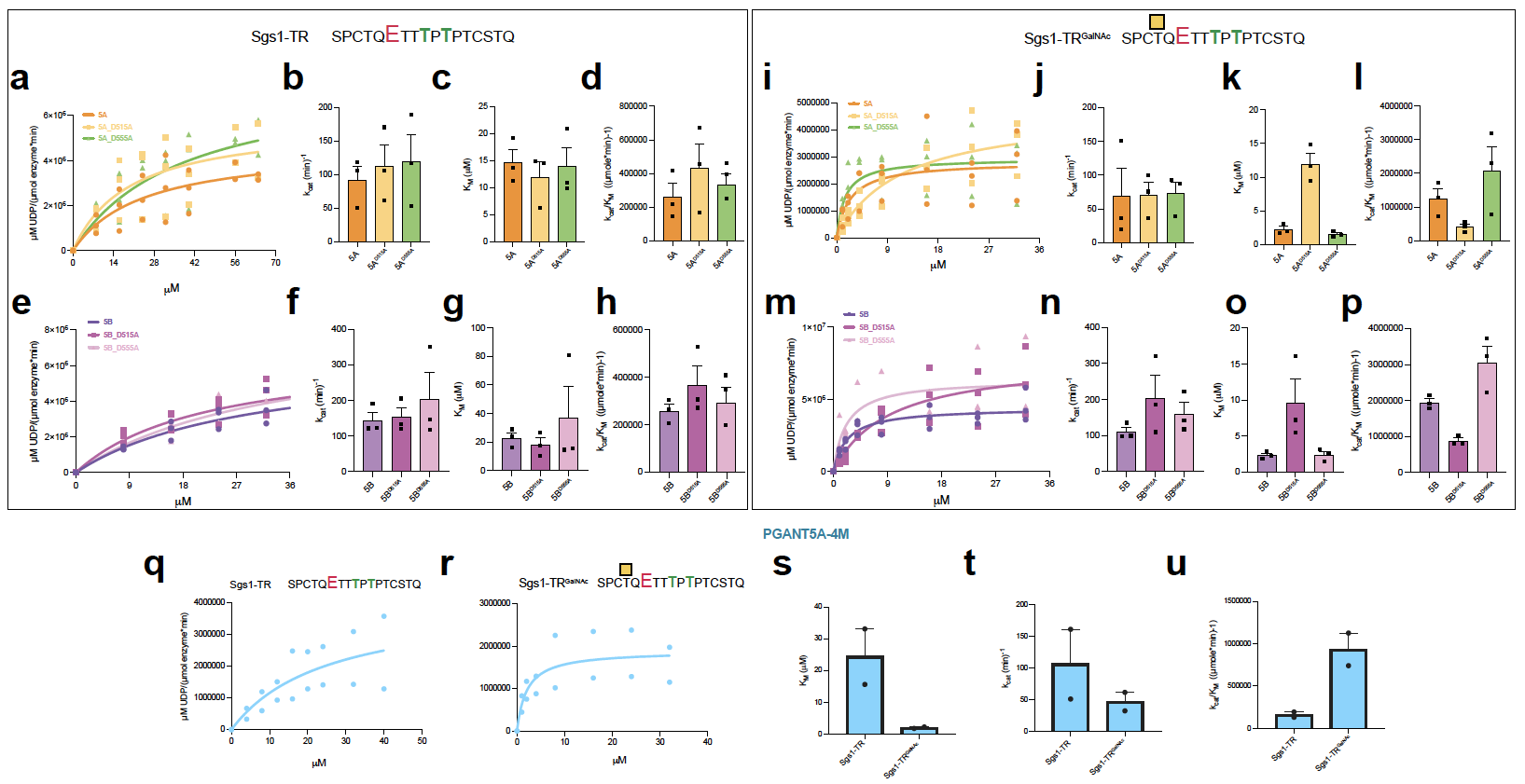

**Supplemental Figure 10. a-h,** Sgs1-TR kinetics for PGANT5A, 5B and lectin domain variants. **i-p,** Sgs1-TR^GalNAc^ kinetics for PGANT5A, 5B and lectin domain variants. **q-u,** Sgs1-TR and Sgs1-TR^GalNAc^ kinetics for PGANT5-4M.

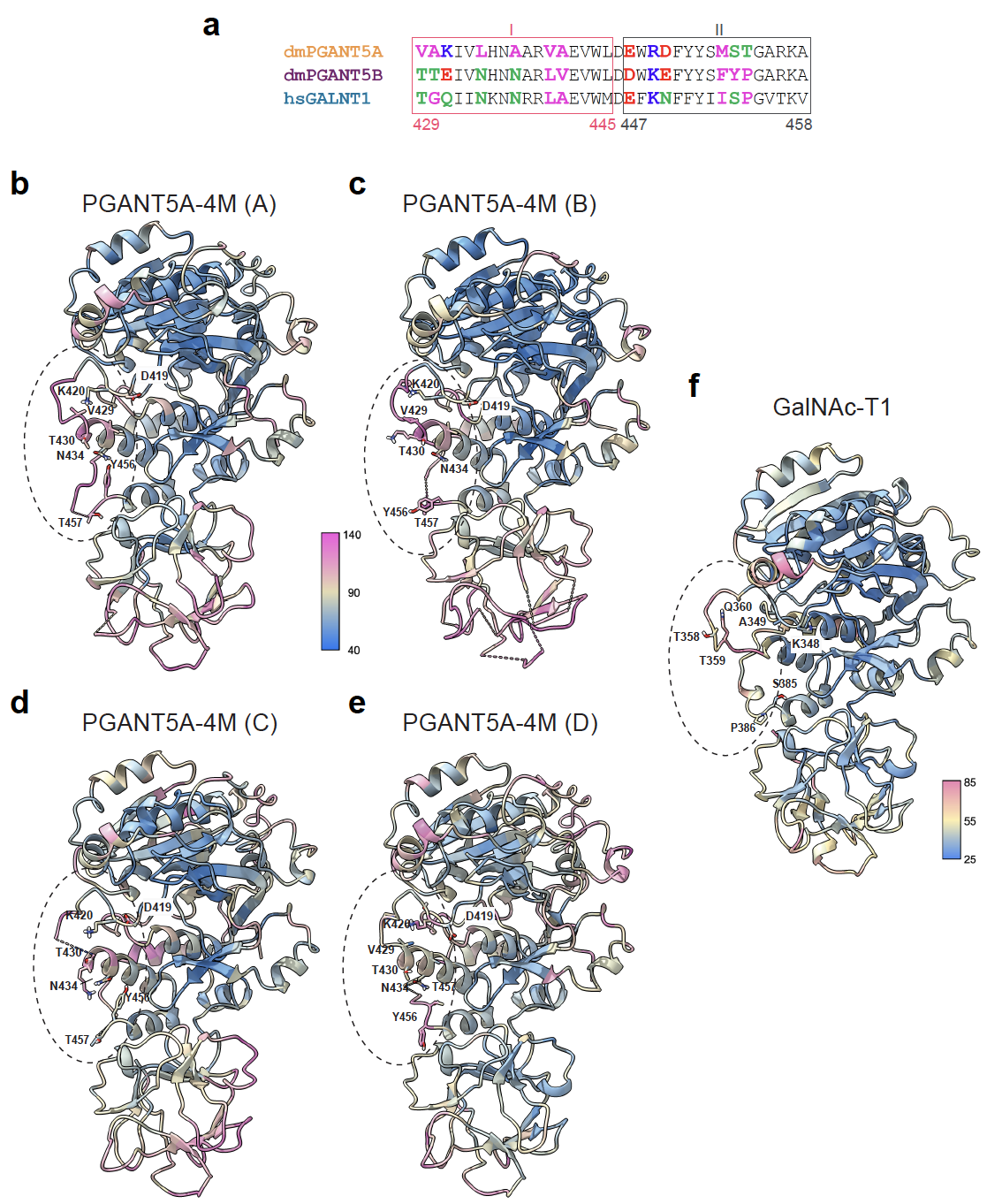

**Supplemental Figure 11. a,** Helix I and II sequence alignments between homologues PGANT5 and GalNAc-T1. AA acid numbering is for PGANT5. **b-e,** B factors mapped onto the crystal structures chains A-D of PGANT5A-4M. **f,** B factors mapped onto the crystal structure of GalNAc-T1 (PDB ID 8V9Q). ChimeraX^1^ was used for B factor mapping and figure generation.

**Supplemental Tables**

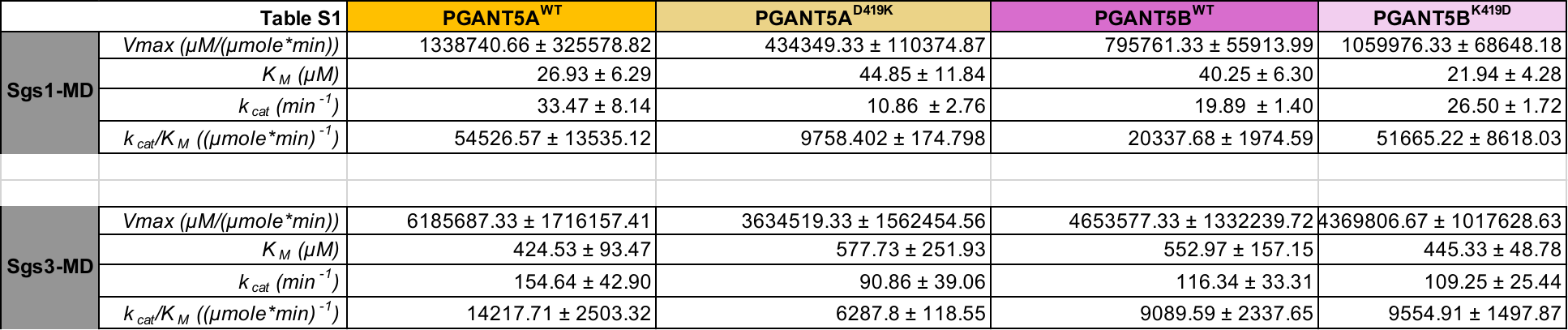

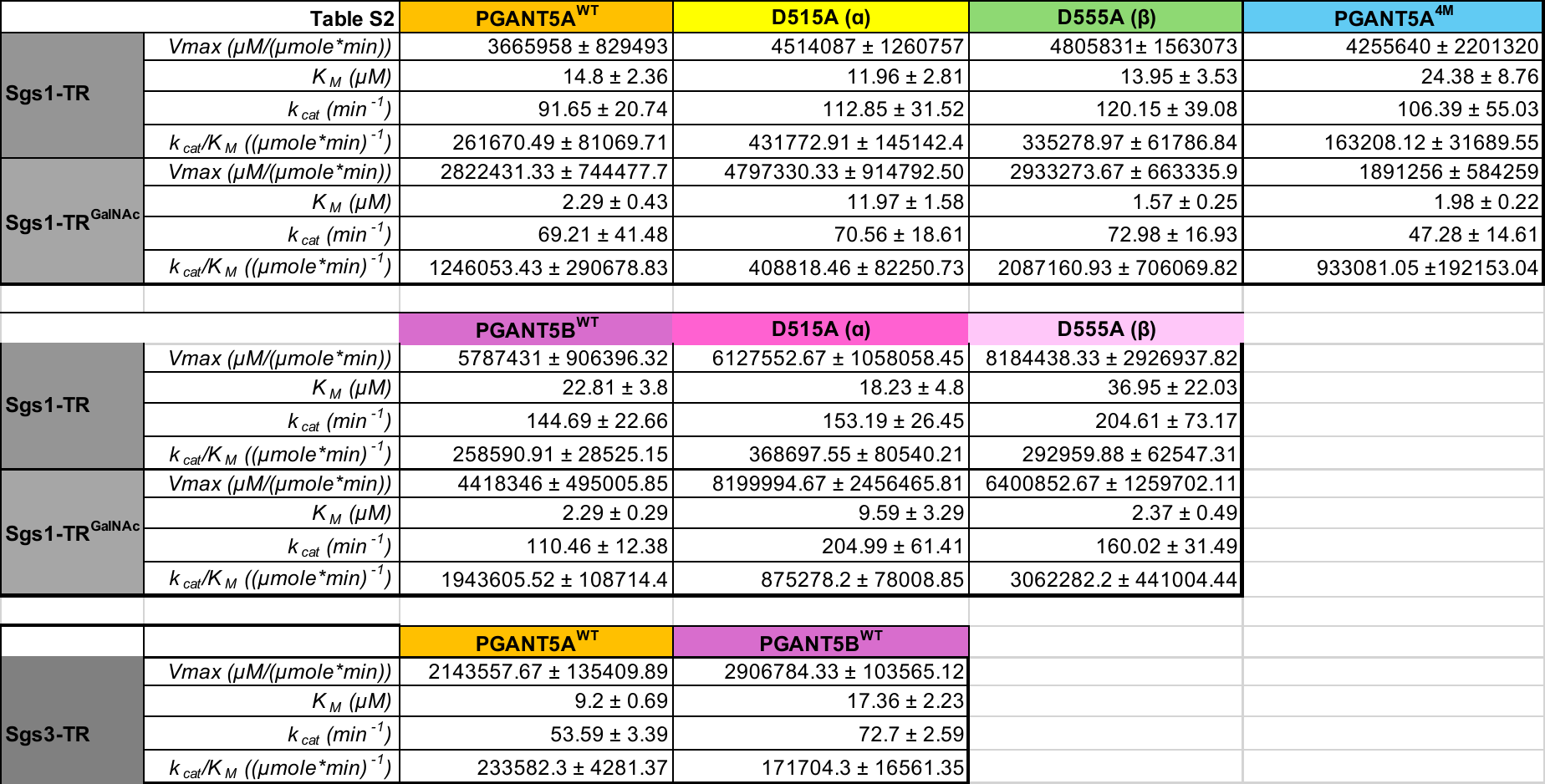

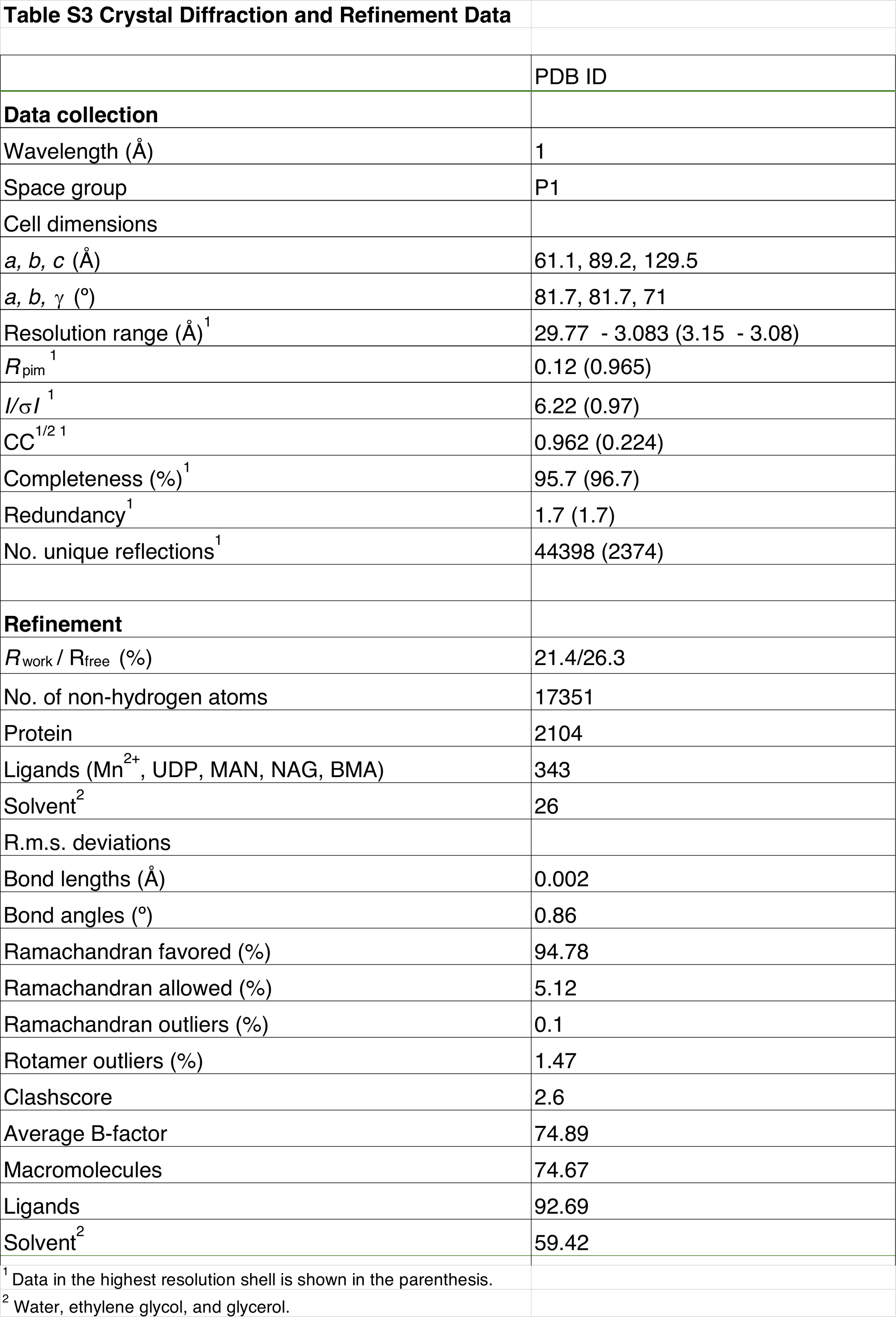
**References**

1	Meng, E. C. *et al.* UCSF ChimeraX: Tools for structure building and analysis. *Protein Sci* **32**, e4792 (2023). <https://doi.org:10.1002/pro.4792>
